# Conserved inhibitory motifs yield distinct hippocampal functions through regional circuit embedding

**DOI:** 10.64898/2026.05.06.718927

**Authors:** Samer Siwani, Angelica Thulin, Arthur S. C. França, Andreas Lindholm, Erika Roman, Klas Kullander

**Affiliations:** Genetics and Neuroscience program, Department of Immunology, Genetics and Pathology, Uppsala University, Uppsala, Sweden; Department of Animal Biosciences, Swedish University of Agricultural Sciences (SLU), Uppsala, Sweden; Netherlands Institute for Neuroscience, RoyalNetherlands Academy of Arts and Sciences, Amsterdam, the Netherlands; Department of Psychiatry, Amsterdam UMC, University of Amsterdam, Amsterdam, the Netherlands; Department of PharmaceuticalBiosciences, Uppsala University, Uppsala, Sweden

**Keywords:** OLM, Approach, Avoidance, Basolateral amygdala, Hippocampus, Emotion, Memory, Nucleus accumbens, Rabies, Tracing

## Abstract

The ventral hippocampus (vHipp) is increasingly recognized for its role in processing probabilistic threats and emotional salience, yet the circuit mechanisms underlying these computations remain unclear. Here, we investigated how oriens-lacunosum moleculare interneurons expressing the α2 subunit of the nicotinic acetylcholine receptor (OLMα2 cells) contribute to hippocampal function along the longitudinal axis. Combining cell-type specific optogenetics, behavioral assays, monosynaptic and transsynaptic viral tracing, we compared intermediate hippocampal (iHipp) and vHipp OLMα2 populations.

We show that OLMα2 cells exert regionally distinct control over behavior. Manipulation of iHipp OLMα2 cells selectively influenced object-directed exploration and novel object processing without affecting anxiety-like behavior, whereas vHipp OLMα2 cells modulated arousal and emotionally driven avoidance without contributing to object memory. Circuit tracing revealed that this functional double dissociation is paralleled by distinct connectivity profiles: iHipp is preferentially embedded in sensory cortical networks, whereas vHipp is strongly connected to limbic structures, including the basolateral amygdala and nucleus accumbens. Notably, despite robust regional amygdala-hippocampal connectivity, OLMα2 interneurons themselves received sparse direct amygdala input, indicating that emotional modulation is mediated indirectly via hippocampal pyramidal cell pathways. In contrast, stimulation of basolateral amygdala projections to vHipp promoted approach behavior without inducing generalized aversion, further supporting pathway- and cell-type specific encoding of valence and salience.

Together, our findings demonstrate that OLMα2 interneurons implement a longitudinally organized inhibitory framework in which conserved microcircuit motifs yield distinct behavioral outcomes, depending on their embedding within region specific input and output networks. This work identifies a mechanistic basis for functional specialization along the hippocampal axis and highlights interneuron positioning as a key determinant of hippocampal contributions to cognition and emotion.

## Introduction

Mounting evidence indicates that the ventral hippocampus (vHipp) is involved in the processing of probable threats, but how this occurs at the circuit level remains unclear. Older studies have assigned the hippocampus as central in the limbic system, however, the currently accepted view considers the hippocampus as a memory related structure that consolidates cortical engrams (Morris et al., 1982; Olton et al., 1979; Scoville and Milner, 1957). Recent studies suggest more immediate processing as well, such as spatial mapping through place and grid cell formations (Moser et al., 2008) and probable threats (Mikulovic et al., 2018; Pi et al., 2020; Wang et al., 2013). The hippocampus is thought to link valences with sensory cues, for example, during fear condition tasks (Phillips and LeDoux, 1992; Sanders et al., 2003). Many studies have focused on the dorsal hippocampus (dHipp), although more recently, investigations of the intermediate (iHipp) and vHipp have been executed (Bast et al., 2009; Burton et al., 2009; Sanders et al., 2003). Differences in gene expression, morphology and connectivity across the hippocampal longitudinal axis have been established (Strange et al., 2014). Projections from structures such as the prefrontal cortex, nucleus accumbens and amygdala are differentiated along this axis (Cholvin et al., 2016; Huff et al., 2016; LeGates et al., 2018; Yang and Wang, 2017), and visual and olfactory inputs project to the dHipp and vHipp respectively (Goodrich-Hunsaker et al., 2008; Hunsaker et al., 2008; Hunsaker and Kesner, 2008; Kesner et al., 2011).

The hippocampus also exhibits a pronounced functional gradient along its longitudinal axis: the dorsal portion is primarily associated with spatial and cognitive processing, while the ventral portion is more strongly linked to emotional regulation and affective behavior (Moser and Moser, 1998; Bannerman et al., 2004; Strange et al., 2014). The iHipp, positioned between these poles, integrates multimodal sensory and contextual information and is thought to mediate the transition between cognitive and emotional domains (Siwani et al., 2018; Komorowski et al., 2013). This contrasts with the dHipp, which does not exhibit comparable responsiveness to emotional stimuli. Notably, activity patterns in the dHipp can even oppose those observed in the iHipp, an effect that may arise from regional differences in cholinergic modulation and nicotine sensitivity (Lovett-Barron et al., 2014; Siwani et al., 2018).

Inhibitory interneurons could contribute to hippocampal subregion-specific functions in at least two non-exclusive ways. First, they might mediate distinct functional roles in each region, meaning that interneurons in the iHipp versus the vHipp implement different circuit operations tailored to the local computational demands; for example, supporting object recognition in the iHipp versus emotional processing in the vHipp (Mikulovic et al., 2018; Siwani et al., 2018). Alternatively, interneurons could operate according to a shared principle of action, providing similar forms of inhibitory control across regions, but producing different outcomes because the afferent inputs and network contexts differ (Lovett-Barron et al., 2012; Klausberger and Somogyi, 2008). In this scenario, the same basic inhibitory mechanism, such as gating excitatory input or regulating timing, underlies region-specific functionality, with the divergence arising from the nature of the incoming sensory or emotional signals rather than intrinsic differences in the interneurons themselves.

Distinct interneuron subtypes, defined by their connectivity, neuromodulatory sensitivity, and molecular markers, regulate the temporal dynamics and input-output relationships of pyramidal cell ensembles (Klausberger and Somogyi, 2008; Lovett-Barron et al., 2012; Dudok et al., 2021). Subtle regional variations in interneuron composition and neuromodulatory control could thus confer specialized modes of information processing across the dorsoventral axis (Jinno and Kosaka, 2006; Andersen et al., 2007; Freund and Katona, 2007). Oriens-lacunosum moleculare interneurons expressing the α2 subunit of the nicotinic acetylcholine receptor (OLM^α2^ cells) are well positioned to regulate the timing and integration of sensory and emotional value signals within hippocampal circuits (Leão et al., 2012; Mikulovic et al., 2018; Siwani et al., 2018). Also, studies of OLM^α2^ cells found that cells positioned in the iHipp are involved in object recognition, whereas OLM^α2^ cells in the vHipp contribute to a behavioral response to predator odor (Mikulovic et al., 2018; Siwani et al., 2018). These findings imply that function is separated depending on the position of OLM^α2^ cells along the intermediate-ventral hippocampal axis. If so, it should be possible to separate their responses and identify whether functionalities might overlap and whether the effects observed are determined by the longitudinal position of the OLM^α2^ populations in the hippocampus. The hippocampus receives differential inputs in the dorso-ventral axis, however, it is not well understood whether these differences apply to the iHipp compared with the vHipp (Amaral and Witter, 1989; Fanselow and Dong, 2010; Risold and Swanson, 1996; Xu et al., 2020; Yang and Wang, 2017).

Thus, OLM^α2^ cells in the iHipp and vHipp may exert either similar or distinct influences on circuit dynamics and behavior, reflecting the differing emotional and motivational demands of the behaviors typically studied—such as avoidance evoked by predator odors versus curiosity-driven exploration of novel objects (Dulawa et al., 1999; Staples, 2010; Toledo-Rodriguez and Sandi, 2011). The vHipp, in particular, has been implicated in both emotional responses and approach behavior toward aversive stimuli (Bland, 1986; Kramis et al., 1975; Montoya et al., 1989; Pi et al., 2020), suggesting that region-specific inhibitory microcircuits may critically shape how emotional and sensory information is processed across the hippocampal axis. To test this hypothesis, we here investigate and compare the connectivity between the iHipp and vHipp, and examine the activity of OLM^α2^ cells in the iHipp and vHipp during behavioral paradigms with distinct emotional and motivational demands, allowing us to assess their role in gating sensory and affective information within hippocampal circuits. In this study we confirm that OLM^α2^ cells in the iHipp and vHipp exert distinct influences on behavior and that this is likely due to the differential communication with external structures.

## Methods

### Experimental animal husbandry

Chrnα2*-cre* and Chrna2-cre/Cas9EGFP transgenic male and female mice (n = 120), with genetic background (Sv129:C57BL/6) as described in our previous study (Leão et al., 2012), at 12-32 weeks of age were used. Mice were kept in an animal room with a 12 hour light/dark cycle (lights on gradually at 07:00-08:00) and constant temperature (22 ± 1 °C) and humidity (50 ± 10%), where they were housed on average 3/cage in 501cm^2^ individually ventilated cages (IVC, GM500). All behavioral procedures were performed during the light cycle. The animals had ad libitum access to food (SSNIFF) and water.

All experimental procedures were in accordance to the approved documentation by Uppsala Animal Ethics Committee (Husbandry permit number: C135/14, 5.8.18-11551.2019, Experimental permit number: C132/13, C45/16 and 5.8.18-07526/2023) and followed the guidelines of the Swedish Legislation on Animal Experimentation (Animal Welfare Act SFS 1998:56 or SFS 2018:1192), the Swedish Animal Welfare Ordinance (SFS 2019:66), the Regulations and General Advice for Laboratory Animals (SJVFS 2019:9, Saknr L 150) and the European Union Directive on the Protection of Animals Used for Scientific Purposes (Directive 2010/63/EU).

### Viral vector injection and optogenetic surgical procedures

Animals were anesthetized using isoflurane, and were given analgesic treatment (Vetergesic/Karprofen and Marcain, Supplementary Table 1) for up to 24 h post-surgery.

For tracing and optogenetic stimulation sub cranial injections of viral vectors containing fluorescent markers and/or opsin containing channels such as archaerhodopsin (Arch) were carried out (Supplementary Table 1) using a nanofil syringe (World Precision Instruments) under a stereotaxic frame, with the following coordinates: Nucleus accumbens: AP: 1.8 mm, ML: −1 mm, DV: −4.3 mm; Basolateral amygdala: AP: −1.7 mm, ML: −2.9 mm, DV: −4.3 mm; Intermediate hippocampus: AP: −3.0 mm, ML: −3.2 and 3.2 mm, DV: −2.7 mm and Ventral hippocampus: AP: −3.2 mm, ML: −3.2 and 3.2 mm, DV: −3.5 mm.

For tracing experiments, virus was injected unilaterally, while for opto-behavioral experiments bilateral injections were performed, followed by implantation of a doric fiber at the most dorsal point of the injection sites, that was secured in place using two anchor screws and dental cement (Intermediate: DV: fiber at −2.7 mm, Ventral: DV: fiber at −3.5 mm, Supplementary Table 1).

### Laser protocols

Green and blue lasers (535 nm and 473 nm, Shangai Dream Lasers, Supplementary Table 1) were used to excite the rhodopsins at ~4 mW. A sinusoidal or flat signal was generated in a DAQ box controlled with a custom Labview interface controlled by EthoVision XT (version 14, 15 and 16; Noldus, Wageningen, the Netherlands) hardware control system. Inhibitory signals were flat in all protocols except the object recognition test to reproduce previous observations (Siwani et al., 2018). In the other tasks a flat laser in a 1-sec interval was used.

### Habituation and handling

Before any behavioral task, mice were first habituated to the experimenter, the experimental room, and the experimental arenas and boxes. Cages were taken to the behavioral room and mice were allowed to acclimatize for 30 min, whereafter the experimenter placed the hands into the cage for 5 min (3 times with >2 hours between sessions). Next, cages were taken to an “intermediary” room and mice were left to acclimatize for 30 min. Then, one by one, a cage was taken into the behavioral room and mice were all placed on the lap of the experimenter where they could explore freely for 5 min (5 times with >2 hours between sessions). Animals that jumped off the experimenter to the floor during handling were excluded (n = 2). During these sessions, light ambience was gradually increased each day from close to 0 lux until it matched the light ambience in the respective behavioral experiment.

The behavioral testing was performed in a succession of mildest to highest stressor. Not all animals went through all the behavioral tasks before post-mortem processing, but the tasks they were tested in, went according to the succession illustrated in Figure 1.

**Figure 1.**
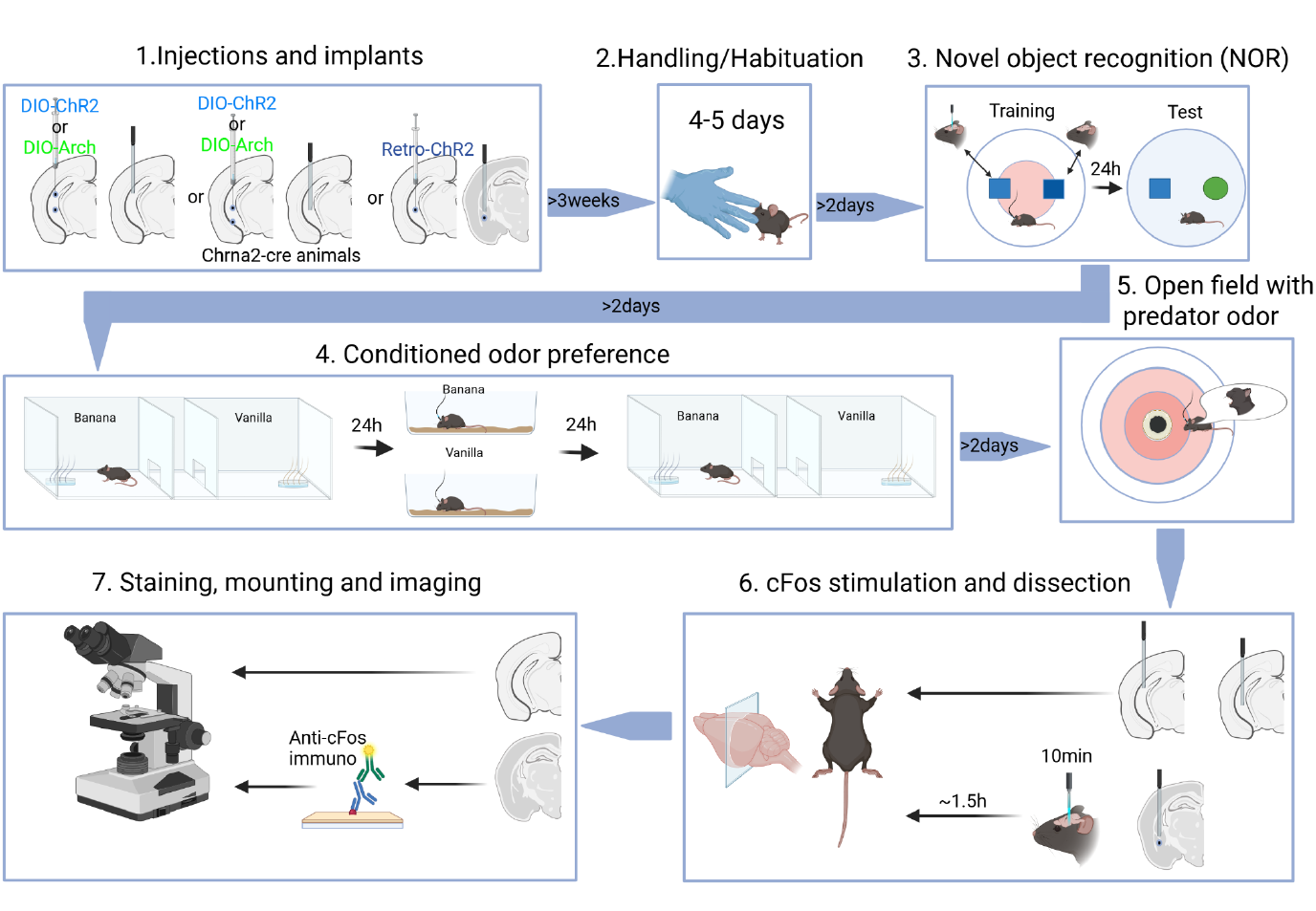
Schematic illustration of the behavior experimental setup and timeline (Created in BioRender. Siwani, S. (2026) https://BioRender.com/d9xh5s0). No individual animal went through all the procedures.

### Novel object recognition (NOR)

The novel object recognition (NOR) test (Figure 1) has been described in detail elsewhere (Siwani et al., 2018). In brief, it consists of a 5 min habituation session to the arena followed by two sessions of 10 min each, 24 hours apart. Two modified brushes were used as objects and were placed in a white plastic circular arena (46 cm in diameter). The objects were placed close enough to the walls so the mice did not need to expose themselves to the center, which is usually avoided, but far enough from the wall for the mouse to pass behind the object without the necessity of exploring it. The behavior of the mice was recorded from above. In the behavioral software EthoVision XT (Noldus, Wageningen, the Netherlands), zones were created around the objects. Size of the zones was determined to be around the head size of the mice from the edge of the object to the edge of the zone. During the training session, two similar objects with slight differences were presented. When the mice approached one of the objects, the laser was turned on (8 Hz). Which object that triggered the laser was randomized between individuals. During the test session, 24 hours later, the objects that did not trigger the laser were switched with a novel object. Time spent (sec) exploring the objects was defined as the amount of time the nose-point was in the zone. Time spent (sec) exploring the center of the arena was defined as the amount of time the center point was in the zone. Distance moved (cm) and velocity (cm/s) was also acquired. Preference ratio in the test session was calculated as time spent in novel object zone/time spent in familiar object zone. The illumination in the arena was measured to be 50-80 lux. In between mice the arena was wiped with 10% ethanol or 70% ethanol followed by water and dried with paper. Any animal that did not explore the objects for more than 10 sec was excluded from the dataset (n = 3). Furthermore, animals that had the optic cables detached (n = 2) and lasers not turning on properly (n = 2) during the training session were excluded. One animal was excluded due to having seizures during the training session.

### Conditioned odor preference (COP)

A three-compartment arena was used for this task (Figure 1), where one “small” (7*15.5 cm) transition compartment was connected to one compartment on each side (15.5*15.5 cm). The behavior of the mice was recorded from above and tracked using EthoVision XT (Noldus, Wageningen, the Netherlands). The illumination in the arena was close to 0 lux. The task consisted of four sessions, one per day:

1. First day, habituation for 10 min. The animal was placed in the middle compartment and allowed to explore the entire arena for 10 min.
2. Second day, preconditioning for 10 min. The mice were kept outside of the room until the setup was prepared. They were then brought in one by one and placed in a resting cage. A cut Q-tip was dipped in a solution of the non-social odors (vanillin ~1 g/100mL in water or isoamyl alcohol 1:100 in water), then placed in a small (3 cm in diameter) petri dish which was positioned in the middle of the wall opposite the door to the transition compartment. The position of each odor (vanillin/isoamyl alcohol) was randomized between the individuals. Q-tips were dipped, and the arena was cleaned (10% ethanol or 70% ethanol followed by water) and dried in between every individual. Duration (sec) and frequency in each compartment were measured.
3. Third day, odor conditioning for 5 min. Two empty home cages were prepared with the two different odors. A Q-tip was dipped in the solution (vanillin/isoamyl alcohol) and was then dragged on the walls of the cage in a zigzag pattern. Then the cue tip was dipped again and was dragged in the sawdust at the bottom of the cage in a zigzag patter. An airlock lid was put on the cage until the task was started. The animals were kept outside of the room until the setup was prepared. The mice were then brought in one by one; fiber optics were connected, and the mouse was placed in a resting cage. Each animal was then placed in each odor cage for 5 min. In one of the odor cages the laser was on the whole time with sinozoidal pulses 10 sec ON, 10 sec OFF for OLM stimulation and 5 sec ON, 10 sec OFF for basolateral amygdala (BLA) stimulation. The odor that was presented first, and the odor that was associated with the laser were randomized across individuals.
4. Fourth day, postconditioning for 10 min. The animals were kept outside of the room until the setup was prepared. They were then brought in one by one and placed in a resting cage. A cut Q-tip was dipped in solution of the non-social odors (vanillin/isoamyl alcohol), then placed in a small (3 cm in diameter) petri dish which was positioned in the middle of the wall opposite the door to the transition compartment. The position of each odor (vanillin/isoamyl alcohol) was randomized between the individuals. Q-tips were dipped, and the arena was cleaned and left to dry in between every individual. Distance moved (cm), velocity (cm/s), duration (sec) and frequency in each compartment were measured.

Preference index was calculated for both time spent and frequency of visits with the odors by the use of the formula below. One outlier was excluded from the OLM-ChR2 dataset using the Grubbs outlier test assuming normal distribution (Figure 7C). One outlier was excluded from the BLA-ChR2 dataset using the ROUT outlier test not assuming normal distribution (Figure 5Q).

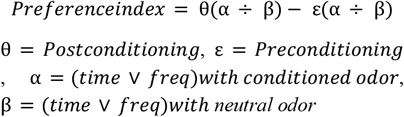

### Predator-odor aversion task (POA)

The POA task was performed in the same arena as the NOR. The procedure has been described in detail elsewhere (Mikulovic et al., 2018). The animals were kept outside of the room until the setup was prepared. They were then brought in one by one; fiber optics were connected, and the mouse was placed in a resting cage. Cat hair from a known hunter house cat was placed in a lidless petri dish with a net mesh placed on top of it. Then this petri dish was placed in the center of the arena (Figure 1). In the behavioral software (EthoVision XT; Noldus, Wageningen, the Netherlands), circular zones were created by dividing the radius into 4 by eye (approx. 4.5 cm each). The center one encircles the petri dish, the next one after that called the intermediate zone and next extended intermediate. The mouse was placed near the wall and time spent (sec) and distance moved with the EthoVision center point in the respective zones was measured. The test was performed in total darkness. The task consisted of a single session of 10 min. The laser was on the whole session with a protocol of square pulses, 1 sec ON and 1 sec OFF in the entire arena. The arena was cleaned between each individual with 10% ethanol and paper towels.

### Postmortem processing

Post behavioral and tracing experiments, mice were given an overdose of anesthetic and analgesic medication mixture (Ketalar and Domitor vet, Supplementary Table 1), before they were perfused with 1xPBS, followed by 4% paraformaldehyde or formaldehyde. The brains were then dissected and fixed in 4% formaldehyde solution overnight. After post-fixation brains were stored in 1xPBS solution until they were sectioned into 70 or 100 µm coronal slices using a vibratome to confirm viral infection and transgene expression. The sections were imaged for expression of marker fluorescent proteins (eYFP, GPF, tdTomato or mCherry; Olympus BX61WI, Zeiss Imager.Z2) using 5x, 10x and 20x objectives under laser illumination and postprocessed in ImageJ or ZEN 3.3 with a stitching tool (Preibisch et al., 2009).

### Immunohistochemistry

#### Anti cFos

Immunohistochemistry against cFos was performed for BLA-ChR2 mice to confirm activation of BLA neurons. Before staining, each animal’s BLA on the right hemisphere was stimulated in an empty cage with light to induce cFos expression. Stimulation was Sinozoidal 8Hz stimulation, 1 sec ON, 1 sec OFF, ~4 mW at the tip for a total of 10 min. ~1.5 hours later the animals were perfused and their brains dissected. 70 µm sections were washed 3×5 min in 1x PBS before incubation in 500 ml blocking solution (1x PBS, 2% Donkey Serum, 1% BSA, 0.1% Triton-X, 0.05% Tween 20, Supplementary Table 1) for 1 h in a 12-well plate and at room temperature (RT). Post blocking the sections were incubated overnight in rabbit anti-cFOS antibody 1:500 dilution in Supermix (1x TBS, 0.25% Gelatine, 0.5% Triton-X) at 4 °C, except for the blocking control that was incubated in the blocking solution. On the second day, sections were washed 3×10 min in TBS (30 g Tris, 80 g NaCl, 2 g KCl in 1000 ml DEPC water, pH 7.4 using HCl to adjust), before they were incubated for 1.5 h at RT in donkey anti-rabbit 647 and 200 nM/ml DAPI in supermix. Thereafter, tissue was washed 4x 10 min in 1x TBST (0.1% Tween20 in 1x TBS) and 1x 10 min in 1x PBS before mounting and imaging. The number of cFos positive cells was counted using the cells counter plugin in ImageJ (Supplementary table 1). Before counting, the BLA borders were identified using Allen Brain Atlas (Allen Reference Atlas - Mouse Brain [brain atlas]. Available from atlas.brain-map.org) and all other structures were removed so as to only count positive cells in BLA. Cells in the right and left BLA were counted in each animal for a comparison across animal groups and left vs right hemisphere within each animal. Animals that lacked expression of the ChR2-tdTomato were excluded (n = 1).

### Anti-GFP and -RFP

70 um sections were washed 3x 5 min in 1x PBS before incubation in 400 ml blocking solution (5% goat serum in 1x PBS containing 0.3% Triton-X) for 2 h at RT in a 12-well plate. Post incubation the sections were incubated overnight at 4°C in Guinea pig anti-GFP or Rabbit anti-RFP antibody (1:1000 dilution in blocking solution, Supplementary Table 1), except for the blocking control that was incubated in blocking solution. During the second day, sections were washed once 3x 10 min in 1x PBS. Then, incubated for 2 h at RT in Alexa 647, Goat anti Guinea pig or Alexa 594, Guinea pig anti Rabbit (1:1000 dilution in 1x PBS containing 0.2% Triton-X and DAPI 1:1000, Supplementary Table 1). After another 3x 10min wash in 1x PBS, sections were mounted and imaged with a Zeiss Imager.Z2 using 5x, 10x and 20x objectives under laser illumination and postprocessed in ImageJ or zeiss microscope software with a stitching tool (Preibisch et al., 2009).

### Tracing

#### General tracing

For these experiments either retrograde tracing AAV containing tdTomato, Chr2-tdTomato or a CamKII driven Arch3.0-GFP containing virus for anterograde tracing were used. (Supplementary Table 1). The viruses were injected in iHipp and vHipp as described above and brains were sectioned into 70 µm sections. The imaged sections were then processed in ImageJ. The expression intensity (from 1-4 where 4 is highest intensity) was determined by the experimenter in comparison to background for each identified structure on all sections. For each animal, every structure’s highest intensity was plotted in a heatmap (Supplementary figure 1). All the observed structures in all the animals are listed and the structures that have not been observed to have expression in the animals receive the grade 0, meaning no expression has been observed (Supplementary Figure 1). For grouped structures we calculated the averages for the expression intensity across the group, only picking structures with expressions that have been observed in the majority of the group. The structures were then grouped in Diencephalon (Hypothalamic and thalamic nuclei), Emotion related (Cingulate and amygdalar areas, bed nucleus of stria terminalis, nucleus accumbens and dorsal peduncular area), Olfactory related, Piriform areas, Prefrontal areas, Rhinal areas, Sensory related (Auditory, somatosensory, visual area), Septum and other (Diagonal band nucleus, agranular insular area, retrosplenial area, caudoputamen, globus pallidus and tanea tecta) and plotted in a heatmap (Figure 4B, J).

### Rabies tracing

For monosynaptic tracing of OLM cells, a cre-dependent TVA containing virus followed by a glycoprotein deleted rabies virus (3 weeks apart) was injected into the iHipp or vHipp (coordinates above) (Supplementary Table 1). For hippocampal projection neuron tracing a retrovirus containing TVA was injected into either Nucleus accumbens (nAcc) or BLA followed by a rabies injection into the ventral hippocampus. Ten days after the rabies injections, the mice were perfused, brains dissected and incubated in 4% FA before sectioning. Anti-RFP and -GFP immunohistochemistry (described above) were done to amplify the signals before imaging. GFP, RFP and eGFP positive cells in the hippocampus (near the injection sites where expression was observed) were quantified. OLM cells were identified based on the morphology, location and expression of eYFP (membrane bound compared to the GFP in TVA positive cells that fills the cells). The number of double positive cells were counted (OLM+rabies and TVA+rabies) and a ratio calculated.

### Statistical analyses

The behavioral raw data were extracted from EthoVision and all data were processed in Microsoft Excel before analyses in GraphPad Prism. The data were checked for normality using the Shapiro-Wilk test before further analysis. Parametric/non-parametric and paired/unpaired T-tests were used to evaluate differences between/within groups or in relation to a theoretical number in each experiment (for instance 0 in the COP task, indicating no shift in preference before and after conditioning) (Supplementary Tables 3, 5 and 6). Two significant outliers were found and removed by applying Grubbs and ROUT outlier tests in two datasets (BLA-ChR2 COP, OLM-ChR2 COP, detected in Figure 5Q and 7C). Data were considered statistically significant at p<0.05.

## Results

### Ventral but not intermediate OLMα2 interneurons modulate emotionally weighted exploration

To investigate how OLMα2 interneurons contribute to behavior across the hippocampal longitudinal axis, we targeted viral opsin expression to either iHipp or vHipp of Chrna2-Cre mice (Figure 2A, B). Expression of eYFP/GFP delivered together with Cre-dependent opsins demonstrated successful delivery of the viral vectors into OLM^α2^ cells in the hippocampal CA1 of Chrna2-Cre mice (Figure 2B). Guided by earlier findings showing that vHipp OLM^α2^ neurons regulate predator-linked behavior (Mikulovic et al., 2018), whereas iHipp OLM^α2^ neurons support object-based memory formation (Siwani et al., 2018), we asked whether manipulating these populations in the *opposite* behavioral domains would reveal latent functional roles.

**Figure 2.**
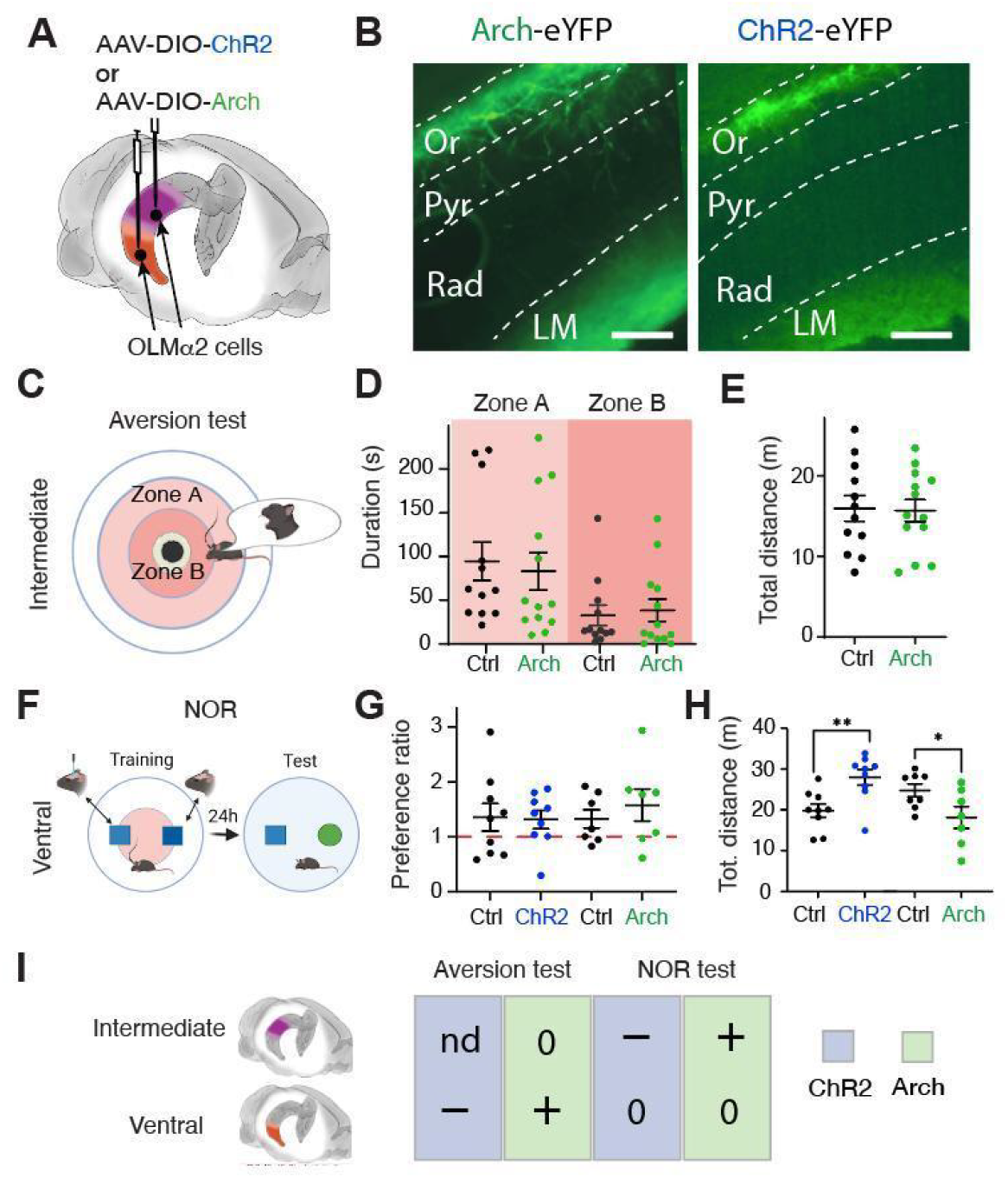
Intermediate and ventral OLM^α2^ cells have differential function in behavior. **A)** Schematic illustration of viruses containing either channelrhodopsin or archaerhodopsin (Char2; Arch; Supplementary Table 1) used for targeting the intermediate (purple) or ventral (orange) hippocampus. **B)** Images demonstrating expression of viral tools in the hippocampus. Scale bars: 100 µm. Abbreviations: LM, lacunosum moleculare, Or, oriens; Pyr, pyramidal; Rad, radiatum. **C)** Schematic of the behavioral setup for the predator odor aversion (POA) test done with modulation of intermediate OLM^α2^ cells, with arena zones indicated (modified from Mikulovic et al., 2018; Created in BioRender. Siwani, S. (2026) https://BioRender.com/d9xh5s0). **D)** The results from the POA test, demonstrating similar duration of time spent irrespective of zones when comparing OLM^α2-Arch^ animals and controls. Lines represent mean ± SEM. Data and statistics are presented in Supplementary Tables 2 and 3. **E)** The results from the POA test, demonstrating similar total distance moved during the task when comparing OLM^α2-Arch^ animals and controls. Lines represent mean ± SEM. Data and statistics are presented in Supplementary Tables 2 and 3. **F)** Schematic illustration of the behavioral setup for the novel object recognition (NOR) test done with modulation of ventral OLM^α2^ cells (Created in BioRender. Siwani, S. (2026) https://BioRender.com/d9xh5s0). **G)** The results for the NOR test, expressed as the preference ratio (time spent with the novel object divided by time with the familiar object), comparing OLM^α2^-manipulated animals (OLM^α2-Arch^ or OLM^α2-ChR2^) with their respective controls. Lines represent mean ± SEM. Data and statistics are presented in Supplementary Tables 2 and 3. **H)** The results for the total distance moved in the NOR test comparing OLM^α2^-manipulated animals (OLM^α2-Arch^ or OLM^α2-ChR2^) with their respective controls. Lines represent mean **±** SEM, *p<0.05, **p<0.0. Data and statistics are presented in Supplementary Tables 2 and 3. **I)** Summary table comparing behavior results from previous studies with the current results (Siwani et al., 2018; Mikulovic et al., 2018). Minus (-) indicates a deficit whereas plus (+) indicates enhancement in NOR performance, or decreased/increased avoidance in the POA task. Zero (0) - no effect, “nd” - not determined.

Following viral expression and implantation of optic fibers (>3 weeks), iHipp-targeted animals were evaluated in a POA test (Figure 2C-E; Supplementary Tables 2 and 3), whereas vHipp-targeted animals were tested in the NOR task (Figure 2F-H; Supplementary Tables 2 and 3). Optogenetic inhibition of iHipp OLM^α2^ cells did not enhance aversion responses, in contrast to the robust effects previously observed for vHipp OLM^α2^ inhibition (Figure 2D, E; Mikulovic et al., 2018; Supplementary Tables 2 and 3). Conversely, manipulation of vHipp OLM^α2^ cells failed to alter object discrimination performance, confirming that these cells do not contribute to NOR, a function instead mediated by iHipp OLM^α2^ cells (Figure 2G, H; Siwani et al., 2018; Supplementary Tables 2 and 3). Interestingly, locomotor activity during the NOR test session was modulated bidirectionally by vHipp OLM^α2^ manipulations: optogenetic activation increased distance moved, whereas inhibition reduced it (Figure 2H; Supplementary Tables 2 and 3). This suggests that vHipp OLM^α2^ cells may influence arousal or exploratory drive without contributing to object memory formation itself. Together, these experiments demonstrate that the behavioral roles of OLM^α2^ interneurons are anatomically partitioned along the hippocampal longitudinal axis (Figure 2I). Ventral OLM^α2^ cells regulate predator-related aversion, whereas intermediate OLM^α2^ cells support object-based memory. Thus, the functional contribution of OLM^α2^ interneurons seem determined by their position within the hippocampal circuit architecture.

### Ventral OLMα2 inhibition induces center avoidance whereas intermediate OLMα2 activation biases object approach

Building on the results above (Figure 2), we hypothesized that the activity changes observed following ventral OLM^α2^ cell modulation might reflect emotion-related alterations during the initial training session, potentially influencing how cautious or exploratory the mice are during subsequent object encounters. To dissect this further, we analyzed behavior specifically during the first exposure to the objects in the NOR training session for both intermediate and ventral groups (Figure 3A, B; Supplementary Tables 2 and 3). Again Chrna2-Cre^+^ and Chrna2-Cre^−^ control animals were injected with Cre-dependent Arch or ChR2 constructs (Figure 3A, Supplementary Table 1). Baseline locomotor activity was unaffected by iHipp or vHipp manipulations (Figure 3C, D, Supplementary Tables 2 and 3), ensuring that subsequent exploration differences were not secondary to motor changes.

**Figure 3.**
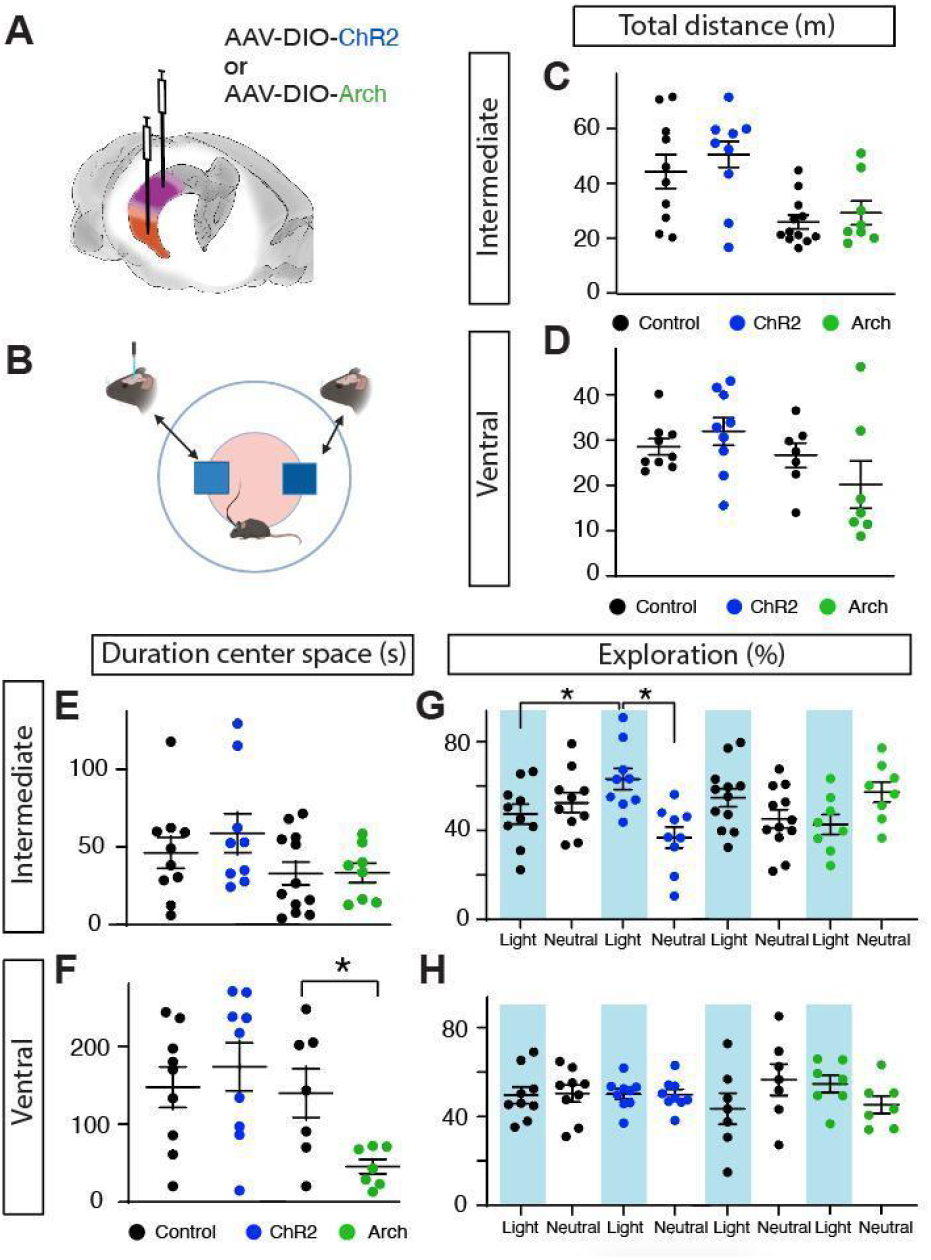
Region-specific modulation of center activity and object preference by OLMα2 cell manipulation. **A)** Schematic illustration of the viral constructs used to target OLM^α2^ cells in the intermediate (purple) or ventral (orange) hippocampus. **B)** Schematic of the novel object recognition (NOR) test showing the arena layout and defined zones (Created in BioRender. Siwani, S. (2026) https://BioRender.com/d9xh5s0). **C-F)** The results for the total distance moved during the NOR training day (**C, D**) and time spent in the center zone (**E, F**) comparing OLM^α2^-manipulated animals (OLM^α2-Arch^ or OLM^α2-ChR2^) with their respective controls with intermediate (**C, E**) and ventral (**D, F**) groups. Lines represent mean **±** SEM. Data and statistics are presented in Supplementary Tables 2 and 3. **G-H)** Object preference expressed as the percentage of time spent exploring a light-paired object (blue background) relative to a neutral object. OLM^α2^-Arch or OLM^α2^-ChR2 manipulations in the vHipp produced no significant differences compared to controls (**H**). In contrast, OLM^α2^-ChR2 stimulation in the iHipp resulted in a significantly higher preference for the light-paired object compared to both non-lit and Cre^-^ controls (**G**). Lines represent mean **±** SEM, *p<0.05. Data and statistics are presented in Supplementary Tables 2 and 3.

We quantified time spent in the center of the arena, investigating animals with iHipp versus vHipp manipulations. Modulating iHipp OLM^α2^ cells did not alter center exploration (Figure 3E; Supplementary Tables 2 and 3). In contrast, inhibition, but not activation, of vHipp OLM^α2^ cells produced a marked avoidance of the arena center (Figure 3F; Supplementary Tables 2 and 3), consistent with a ventral hippocampal role in anxiety-related behavioral states.

We next examined object approach behavior by comparing exploration of the light-paired object versus the neutral object (Figure 3G, H). Stimulation of iHipp OLM^α2^ cells promoted preferential exploration of the light-paired object, whereas inhibition had no effect (Figure 3G; Supplementary Tables 2 and 3). Notably, this initial bias did not yield stable memory, as these animals later failed to discriminate the familiar object in the NOR test during the test phase (Figure 2I), indicating that enhanced approach does not translate into long-term object encoding. Ventral OLM^α2^ manipulation did not influence object approach (Figure 3H; Supplementary Tables 2 and 3), further supporting the view that these neurons affect emotional bias rather than object salience per se.

### Intermediate and ventral hippocampus exhibit differential connectivity with external structures

We hypothesized that the distinct behavioral phenotypes observed for iHipp and vHipp OLM^α2^ cells could reflect differences in their long-range connectivity. To test this, we injected iHipp and vHipp regions with a viral vector under a general promoter designed to infect fibers at the injection site and label distal projection neurons (Figure 4A-H; Tervo et al., 2016). Injections were made at the same coordinates used for the behavioral experiments, allowing for comparison between connectivity and function (Figure 4C, F). Following expression, projection labeling intensity was scored qualitatively on a 5-step scale (Figure 4D, E, G, H, Supplementary Figure 1), and the results were summarized in a heatmap to visualize the relative connectivity differences between the two regions (Figure 4B). Analysis revealed pronounced differences in the pattern and strength of projections to external structures, highlighting that iHipp and vHipp are embedded within largely distinct extrahippocampal networks (Figure 4B). However, we observed that the virus exhibited mixed anterograde and retrograde labeling tendencies, potentially complicating interpretation. To address this, we complemented the analysis with additional tracing approaches to more specifically delineate inputs and outputs of the two regions.

**Figure 4.**
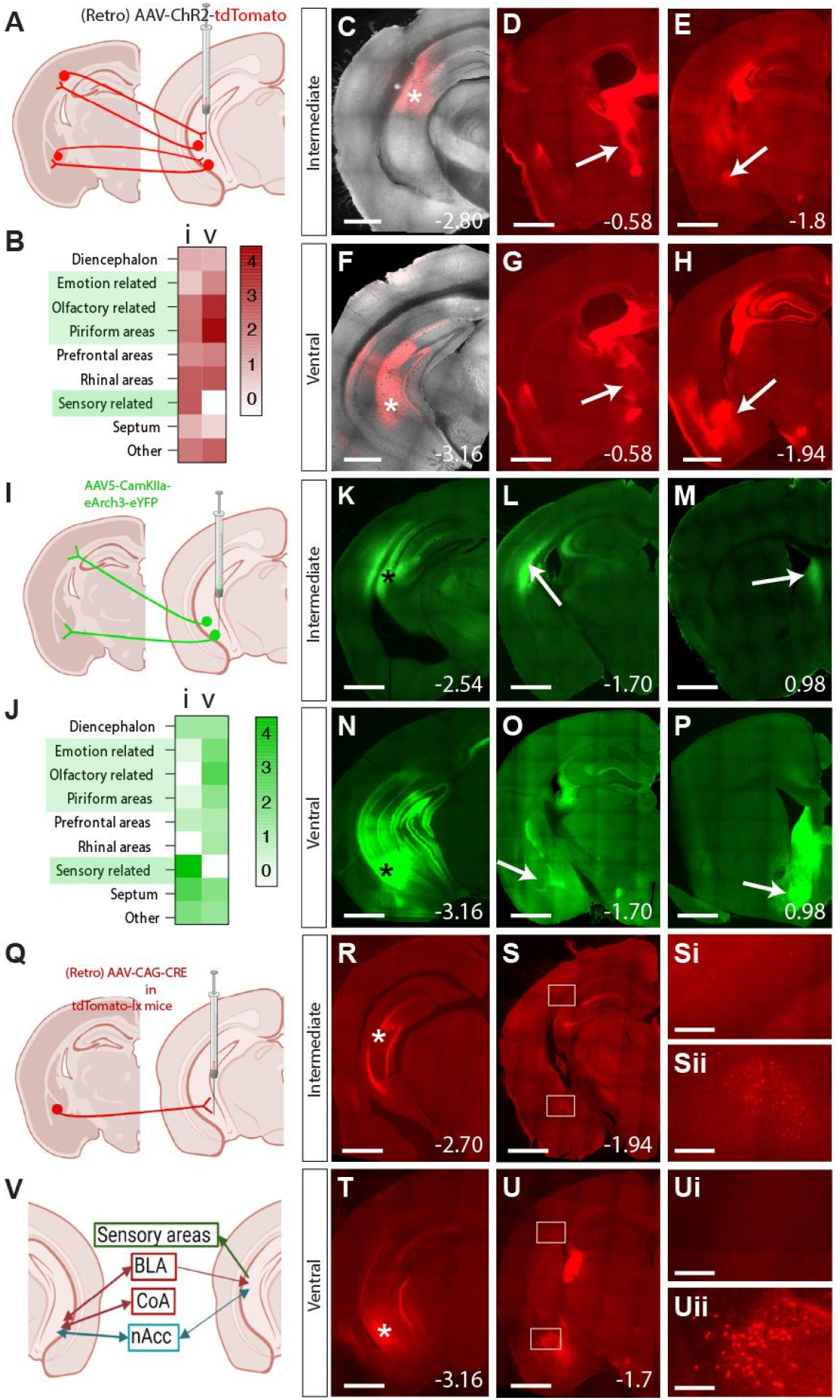
Intermediate and ventral hippocampus display different patterns of connectivity with external structures. **A)** Schematic illustration of the tracing procedure with the rAAV-CAG-hChR2(H134R)-tdTomato virus, in this setting used as a general tracer (Created in BioRender. Siwani, S. (2026) https://BioRender.com/3si8dgl). **B)** Color-coded heatmap summarizing projection strength across grouped brain regions. Comparisons are shown for iHipp (i, left) and vHipp (v, right), with expression intensity represented on a five-point scale (0-4). Regions most relevant to the behavioral effects are shown with a green background. **C, F)** Images of coronal brain sections showing viral injection sites (white star) and spread as indicated by expression of tdTomato red fluorescence in the iHipp (**C**) and vHipp (**F**). **D, E, G, H)** Images of coronal brain sections showing tdTomato fluorescence following tracing using rAAV-CAG-hChR2(H134R)-tdTomato. Injections into iHipp (**D, E**) and vHipp (**G, H**) reveal differential projection patterns; arrows point to thalamic (**D, G**) and amygdalar (**E, H**) regions. **I)** Schematic illustration of the tracing procedure with the AAV5-CAmKIIa-eArch3-EGFP virus used as an anterograde tracer (Created in BioRender. Siwani, S. (2026) https://BioRender.com/3si8dgl). **J)** Color-coded heatmap summarizing projection strength across grouped brain regions. Comparisons are shown for iHipp (i, left) and vHipp (v, right), with expression intensity represented on a five-point scale (0-4). Regions most relevant to the behavioral effects are shown with a green background. **K, N)** Images of coronal brain sections showing viral injection sites (black star) and spread as indicated by expression of green fluorescence in the iHipp (**K**) and vHipp (**N**). **L, M, O, P)** Images of coronal brain sections showing GFP fluorescence following anterograde tracing and signal amplification using an anti-GFP antibody. Injections into iHipp (**L, M**) and vHipp (**O, P**) reveal differential projection patterns, indicated by arrows pointing to sensory (**L, M**) and emotion (**O, P**) related regions. **Q)** Schematic illustration of the tracing procedure with the rAAV-CAG-CRE virus used as a retrograde tracer (Created in BioRender. Siwani, S. (2026) https://BioRender.com/3si8dgl). **R, T)** Images of coronal brain sections showing viral injection sites (white star) and spread as indicated by expression of tdTomato red fluorescence in the iHipp (**R**) and vHipp (**T**). **S, U)** Images of coronal brain sections showing tdTomato fluorescence following retrograde tracing. Injections into iHipp (**S**) and vHipp (**U**) show absence of expression in somatosensory cortex (**Si, Ui**) and differential expression in the amygdala (**Sii, Uii**). **V)** Schematic illustration of summed differences comparing tracing to and from the intermediate and ventral hippocampus (Created in BioRender. Siwani, S. (2026) https://BioRender.com/3si8dgl). Scale bars: 500 µm (except Si,ii and Ui,ii: 100 µm). Distance from bregma as indicated in accordance with Paxinos et al., 2001.

For output mapping, we injected a CamKII-driven Arch-GFP viral vector into iHipp or vHipp (Figure 4I-P). Selected sections across the anterior-posterior axis were amplified with anti-GFP immunostaining to visualize axonal projections. This approach revealed pronounced differences in hippocampal outputs: iHipp projected prominently to cortical sensory areas, including auditory, somatosensory, and visual regions (Figure 4J, L, M, Supplementary Figure 1), whereas vHipp exhibited stronger projections to emotion-related structures such as the amygdala (Figure 4J, O, P Supplementary Figure 1). These results suggest that intermediate and ventral hippocampal regions are embedded in largely distinct cortical and subcortical networks, consistent with their divergent roles in exploratory and affective behaviors.

To examine afferent connectivity, we used a retrograde viral approach in tdTomato-lx mice to label neurons projecting to either iHipp or vHipp (Figure 4Q-U). This allowed us to test whether the basolateral amygdala (BLA) and cortical sensory areas provide input to these hippocampal regions. We observed no projections from cortical sensory areas to either iHipp or vHipp (Figure 4S, Si, U, Ui), whereas the BLA projections were detected in both regions (Figure 4Sii, Uii), with a pronounced BLA projection to the vHipp (Figure 4Uii).

Together, these complementary anterograde and retrograde tracing experiments indicate that iHipp and vHipp differ in both their inputs and outputs. The iHipp is preferentially connected to sensory cortical areas, whereas the vHipp is more strongly embedded in emotion-related circuits, particularly the amygdala (Figure 4V). These anatomical distinctions provide a structural framework that may underlie the region-specific behavioral roles observed for OLM^α2^ interneurons.

### Stimulation of BLA projections to vHipp induces object preference without affecting negative valence behaviors

To investigate whether direct activation of BLA neurons projecting to vHipp could also modulate behavior, we injected a retrograde ChR2-tdTomato virus or tdTomato control virus into the vHipp (Figure 5A) and implanted an optical fiber above the BLA (Figure 5B). After the behavioral tasks, optical stimulation was delivered unilaterally to the BLA in the right hemisphere, and cFos expression was analyzed to verify effective activation (Figure 5C-H). Comparison of left and right BLA revealed a significant increase in cFos-positive cells in the stimulated BLA of ChR2-expressing mice (Figure 5E; Supplementary Tables 4 and 5), with a significant difference relative to control animals in right-left cFos activity (Figure 5H; Supplementary Tables 4 and 5). These results confirm that optical stimulation reliably activated BLA neurons projecting to vHipp.

**Figure 5.**
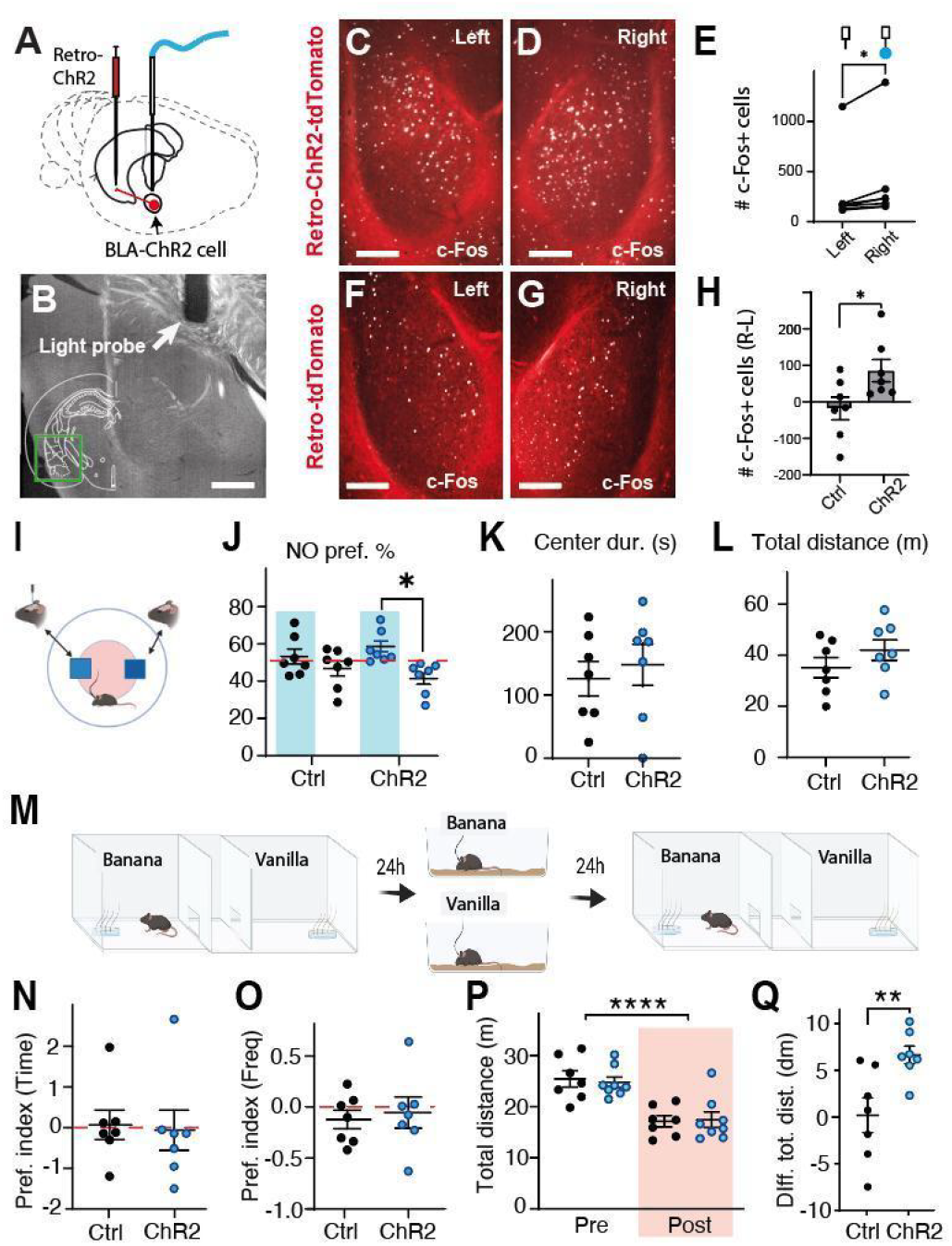
Object preference was induced by stimulation of BLA projection neurons targeting the hippocampus. **A)** Schematic of the experimental design showing viral injection into the basolateral amygdala (BLA) and positioning of the optical fiber cannula in the hippocampus. **B)** Post experimental image showing the placement of the optical fiber cannula (light probe). Green square denotes areas shown in **C, D, F, G**. Scale bar: 100 µm**C, D, F, G)** Representative expression patterns of tdTomato in left and right BLA in control animals (**F, G**) and ChR2-tdTomato in stimulated animals (**C, D**). cFos immunostaining (white) was used to confirm BLA activation following light stimulation of right BLA. Scale bar: 100 µm **E, H)** Quantification of cFos activity in BLA comparing the right (stimulated) and left (non-stimulated) side in ChR2+ mice (**E**) and difference in cell activity (right-left) between control and ChR2+ animals (**H**). Data and statistics are presented in Supplementary Tables 4 and 5. **I)** Schematic of the single trial NOR task with a lit object (left square), non-lit object (right square) and a center zone (pink circle (Created in BioRender. Siwani, S. (2026) https://BioRender.com/d9xh5s0). **J-L)** Behavioral results showing object preference in exploration % (**J**), time spent in the center zone (**K**), and the total distance moved (**L**) during the NOR task comparing BLA-ChR2 to controls. Lines represent mean **±** SEM. Data and statistics are presented in Supplementary Tables 4 and 5. *p<0.05. There is a significant preference in the BLA-ChR2+ animals for the light-paired object **M)** Schematic of the four-day conditioned odor preference (COP) task (Created in BioRender. Siwani, S. (2026) https://BioRender.com/d9xh5s0). **N-P)** Behavioral results from the COP test showing the odor preference index (conditioned odor vs other odor) before and after conditioning, quantified as time spent (**N**) and frequency of visits (**O**), as well as the total distance moved during pre- and post-conditioning days (**P**) during the odor task trials comparing BLA-ChR2 and controls. Lines represent mean **±** SEM. Data and statistics are presented in Supplementary Tables 4 and 5. **Q)** Results from the COP test showing the total distance moved during the odor task conditioning day subtracting conditioned odor from the neutral odor. ChR2+ animals showed an increased activity with conditioned odor compared to controls. Lines represent mean **±** SEM. Data and statistics are presented in Supplementary Tables 4 and 5.

We next assessed behavioral effects in an object exploration task analogous to that used previously for OLM^α2^ cells (Figure 5I). Stimulation of BLA projections to vHipp induced a robust preference for the light-paired object (Figure 5J, Supplementary Tables 4 and 5). Notably, there was no effect on anxiety-related measures such as center duration (Figure 5K, Supplementary Tables 4 and 5) or general locomotor activity (Figure 5L, Supplementary Tables 4 and 5), suggesting that BLA→vHipp stimulation promotes selective approach behavior without eliciting generalized negative-valence responses. Although this may appear contradictory to the role of vHipp OLM^α2^ cells in emotionality observed in other tasks, it likely reflects the engagement of distinct pyramidal neuron populations that mediate positive or exploratory behaviors rather than OLM^α2^-mediated avoidance circuits.

To further probe potential effects on emotional arousal, we adapted the conditioned place preference paradigm by replacing visual cues with neutral olfactory cues, creating a conditioned odor preference (COP) task (Figure 4M; Crawley et al., 2007; Wersinger et al., 2007; Woodley and Baum, 2003; Wrenn et al., 2004; Yang and Crawley, 2009). This allowed us to test whether modulation of BLA→vHipp projections could induce negative-valence associations, given that OLM^α2^ cells play a crucial role in predator odor responses (Mikulovic et al., 2018). No significant changes in odor preference were observed between pre- and post-conditioning in ChR2-BLA or control animals (Figure 5N, O; Supplementary Tables 4 and 5). Likewise, general activity levels did not differ between groups, with only within-group significant differences observed from pre-to post-conditioning (Figure 4P; Supplementary Tables 4 and 6). However, during the conditioning day in the odor task, we noted that BLA-ChR2 mice seemed to display increased activity. To quantify this, we analyzed general activity levels during conditioning and found that stimulation of BLA→vHipp projections increased overall activity (Figure 4Q, Supplementary Tables 4 and 5). These findings indicate that selective activation of BLA projections to vHipp promotes object-directed preference without eliciting aversion, consistent with a role for this pathway in modulating approach behaviors rather than negative-valence responses.

### Monosynaptic tracing demonstrates sparse labelling of amygdala neurons projecting to the ventral and intermediate OLM^α2^ interneurons

Based on our connectivity data (Figure 4), which showed that the majority of BLA projections target the vHipp, we focused on vHipp OLM^α2^ cells to examine whether they receive direct amygdala input. To address this, we performed monosynaptic retrograde rabies tracing in combination with Cre-dependent TVA-GFP expression. TVA-GFP was injected into the vHipp of Chrna2-Cre mice, followed by glycoprotein-deleted rabies virus expressing mCherry (Figure 6A). This revealed sparse labeled projection neurons in the posterior amygdaloid nucleus (PA) connecting to vHipp OLM^α2^ cells (Figure 6C, Ci), whereas labeling within the amygdala was not observed. These data indicate that the behavioral effects observed following vHipp OLM^α2^ manipulation are unlikely to result from direct BLA input. We therefore hypothesized that vHipp OLM^α2^ cells modulate behavior indirectly via hippocampal pyramidal neurons that project to emotion-related structures. To test this, we injected a retrograde TVA-expressing virus into either the BLA or nAcc, combined with rabies virus injection into vHipp (Figure 6D, G). This allowed identification of hippocampal projection neurons that are postsynaptic targets of OLM^α2^ interneurons and communicate with these downstream structures. We observed that vHipp OLM^α2^ cells connect to pyramidal neurons projecting to both BLA and nAcc (Figure 6F, Fi and I, Ii), with similar proportions of OLM^α2^/rabies-labeled cells relative to TVA/rabies double-positive pyramidal neurons in both pathways (Figure 6Fii, Iii). Notably, rabies positive cells were found in the BLA, likely originating from BLA-vHipp projections that had taken up rabies virus particles via their axon terminal in the vHipp (Figure 6E, Ei). Such rabies positive cells were however not found in the nAcc (Figure 6H, Hi), suggesting that vHipp projection neurons that innervate the BLA receive feedback from neurons in the BLA, but not from neurons in the nAcc.

**Figure 6.**
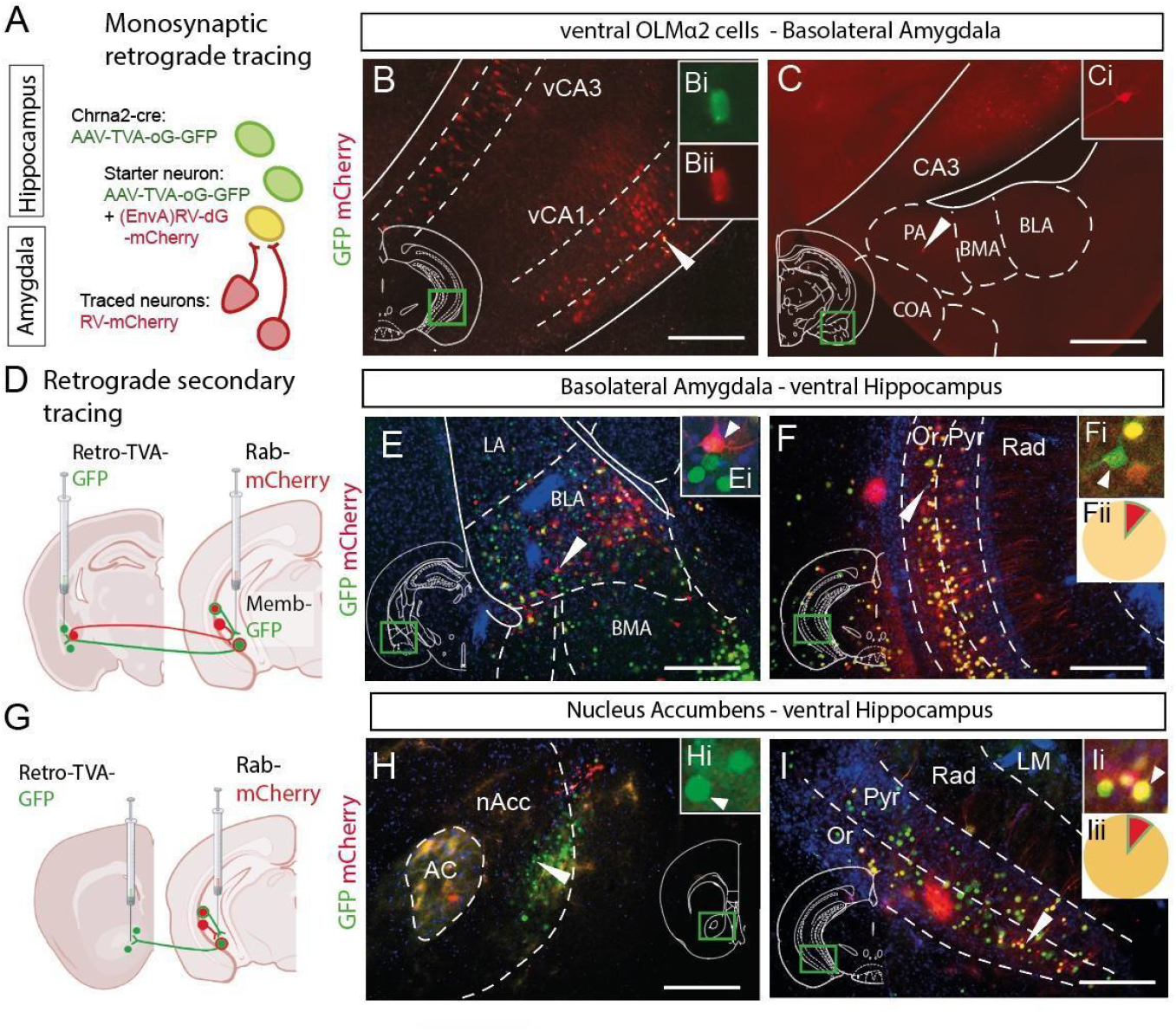
Monosynaptic tracing demonstrates sparse labelling of amygdala neurons projecting to the ventral and intermediate OLM^α2^ interneurons. **A)** Schematic of the rabies tracing showing the targeted OLM^α2^ interneurons and labeled connections. **B)** Expression of labeled cells near ventral hippocampus injection site with avian receptor protein (TVA) expression reported by GFP (green) and dG-Rabies by mCherry (red). **Bi, ii)** Close-up of example starter cell (arrow in **B**), red (**Bii**) and green (**Bi**) channel separated. Scale bar: 150 µm. Abbreviations: vCA1, ventral cornu ammonis area 1; vCA3, ventral cornu ammonis area 3. **C)** Image of the amygdala region with one RV-mCherry labeled cell found in the posterior nucleus of the amygdala (PA). **Ci)** Close-up. Scale bar: 150 µm. Abbreviations: CA3, cornu ammonis area 3; COA, cortical amygdala; BLA, basolateral amygdala; BMA, basomedial amygdala. **D, G)** Schematics of imaged regions and projection neurons labeled in the tracing experiment with either retro-TVA-GFP virus injected into BLA (**D**) or nAcc (**G**; Created in BioRender. Siwani, S. (2026) https://BioRender.com/3si8dgl). **E, H)** Images of neurons labeled by the rabies tracing of hippocampal projections to BLA (**E**) and nAcc (**H**). **Ei, Hi)** close up of cells expressing Retro-TVA-GFP (green; **Hi**) and/or RV-mCherry (red; **Ei**). Scale bars: 150 µm. Abbreviations: AC, anterior commissure; BLA, basolateral amygdala; BMA, basomedial amygdala; LA, lateral amygdala; nAcc, nucleus accumbens. **F, I)** Images of neurons labeled by the rabies tracing of hippocampal projections to BLA (**F**) and nAcc (**I**) with close up of examples (**Fi, Ii**). Pie charts indicate the percentages of double positive cells that are identified as TVA positive projection neurons to BLA (**Fii**, 88.4% in yellow) vs OLM^α2^ cells (**Fii**, 11.6% in red), whereas TVA positive projection neurons to nAcc (**Iii**, 88.7% in yellow) vs OLM^α2^ cells (**Iii**, 11.3% in red). Scale bars: 150 µm. Abbreviations: LM, lacunosum moleculare; Or, oriens; Pyr, pyramidal; Rad, radiatum

Together, these results indicate that ventral OLM^α2^ interneurons are primarily targeted for their potential role in emotion-related behaviors, and that they exert their effects indirectly by modulating hippocampal projection neurons that communicate with downstream affective circuits rather than receiving strong direct input from the BLA.

### Stimulation of ventral OLMα2 cells enhances preference for a conditioned odor

To determine whether modulation of vHipp OLM^α2^ cells influences arousal-related behavior, we expressed either ChR2 or Arch (Figure 7A) in these neurons and tested animals in the COP task. We observed no significant shift in total time spent in the odor chamber for either Arch or ChR2 groups (Figure 7B; Supplementary Tables 4 and 5). However, ChR2 animals exhibited a significant increase in the frequency of visits to the conditioned odor chamber, an effect not observed in the Arch group (Figure 7C; Supplementary Tables 4 and 5). General locomotor activity did not differ between groups, with significant changes observed only within groups from pre-to post-conditioning (Figure 7D, E; Supplementary Tables 4 and 5).

When we analyzed general activity levels during conditioning, we found that stimulation of BLA→vHipp projections increased overall activity (Figure 5Q), however, that was not found upon stimulation of ventral OLM^α2^ cells (Figure 7F, Supplementary Table 4 and 5). These results suggest that ventral OLM^α2^ cell activation enhances approach-related aspects of conditioned odor exploration without broadly increasing arousal, whereas BLA activation induces more generalized activity changes consistent with heightened emotional reactivity.

**Figure 7.**
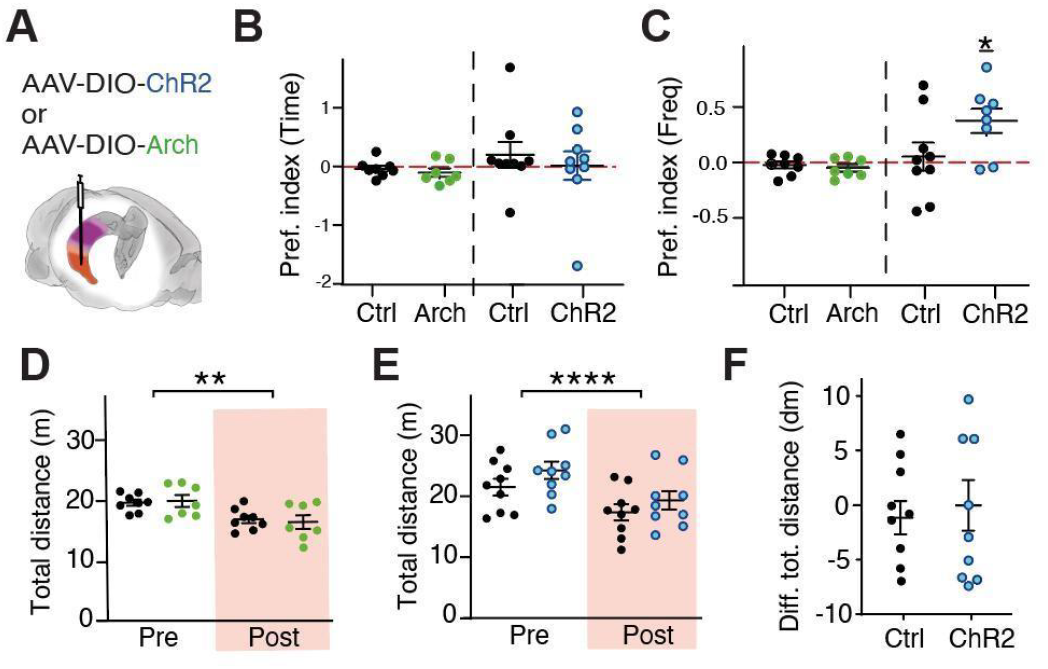
Conditioning of a neutral odor by stimulation of ventral OLM neurons. **A)** Schematic illustration of the viral constructs used to target OLM^α2^ cells in the ventral (orange) hippocampus. **B, C)** Behavioral results showing the odor preference index, calculated from the time spent in the odor chambers (**B**) or the frequency of visits to the odor chambers (**C**), comparing OLM^α2^-Arch and OLM^α2^-ChR2 animals with their respective controls. Lines represent mean ± SEM. Data and statistics are presented in Supplementary Tables 4 and 5. *p<0.05. **D, E)** Results from behavioral experiments showing the total distance moved during the odor task, comparing OLM^α2^-Arch (**D**) and OLM^α2^-ChR2(**E**) animals with their respective controls. Lines represent mean ± SEM. Data and statistics are presented in Supplementary Tables 4 and 6. **p<0.01, ****p<0.0001. **F)** Total distance moved during the odor task conditioning day comparing conditioned odor and other odor (conditioned odor-other odor). Lines represent mean ± SEM. Data and statistics are presented in Supplementary Tables 4 and 5.

## Discussion

Our findings demonstrate that OLM^α2^ interneurons along the longitudinal hippocampal axis regulate distinct behavioral domains through region-specific circuit architectures. By combining precise optogenetic manipulations with complementary anterograde, retrograde, and monosynaptic tracing, we show that intermediate hippocampal (iHipp) OLM^α2^ cells contribute primarily to object-based memory, whereas ventral hippocampal (vHipp) OLM^α2^ cells modulate affective and arousal-related behaviors. These results provide a mechanistic explanation for long-standing observations that hippocampal computations vary systematically along the septotemporal axis (Fanselow and Dong, 2010; Strange et al., 2014; Lisman and Redish, 2009) and highlight interneuron subtype-specific contributions to this functional gradient.

### Circuit organization underlies the functional segregation of OLM^α2^ interneurons

Decades of anatomical work has established that the vHipp is tightly embedded within limbic networks, receiving dense input from and projecting to regions such as the BLA, prefrontal cortex, and nAcc (Swanson and Cowan, 1977; Pitkänen et al., 2000; Strange et al., 2014). In contrast, the iHipp occupies a transitional zone enriched in sensory-related cortical inputs and outputs (Ito et al., 2015; Dollár et al., 2022). Our connectivity analyses, using viral tracers, CamKII-driven anterograde labeling, and retrograde tdTomato reporters, confirmed and extended these observations: iHipp displays prominent projections to sensory cortices, whereas vHipp exhibits stronger connectivity with affective structures, especially the amygdala (Figure 4 and Supplementary Figure 1).

A recent tracing study provides complementary evidence at the level of interneuron inputs. Using monosynaptic rabies tracing, the authors show that OLM^α2^ cells receive markedly different presynaptic inputs depending on their position along the dorsoventral axis (Thulin et al., 2025) These anatomical distinctions create different computational contexts for OLM^α2^ interneurons. Because OLM cells regulate the integration of entorhinal and CA3 inputs onto CA1 pyramidal neurons (Leão et al., 2012; Mikulovic et al., 2015; Malerba et al., 2016), their functional output depends on the downstream targets of these pyramidal neurons. Our tracing experiments showed that vHipp OLM^α2^ cells modulate pyramidal neurons projecting to both BLA and nAcc, forming a disynaptic route that allows these interneurons to shape limbic computations indirectly (Figure 6). In contrast, projection targets of iHipp pyramidal neurons include sensory cortices and multimodal associative areas, providing a substrate for OLM^α2^ regulation of object encoding and perceptual integration during memory tasks (Figure 4 and Supplementary Figure 1).

### Ventral OLM^α2^ interneurons modulate emotional/arousal circuits without receiving strong direct amygdala input

A key mechanistic insight from our study is that, despite strong BLA→vHipp projections at the regional level, monosynaptic rabies tracing revealed surprisingly sparse direct input from amygdala neurons to OLM^α2^ cells (Figure 6C). This indicates that OLM^α2^ modulation of affective behavior does not arise from direct amygdaloid drive. Instead, our experiments demonstrate that vHipp OLM^α2^ interneurons act primarily through local CA1 microcircuits, controlling the excitability and output of pyramidal neurons that themselves communicate with affective centers.

This finding aligns with work showing that hippocampal interneurons often influence behavior by gating pyramidal cell ensembles rather than by receiving strong long-range synaptic drive (Lovett-Barron et al., 2014; Lovett-Barron et al., 2020; Xu et al., 2016). The selective feedback observed from BLA to vHipp projection neurons, but not to OLM^α2^ cells, supports a model in which affective information re-enters the hippocampus through pyramidal-cell pathways, enabling recurrent loops that can be modulated by local inhibitory signals.

### Divergent behavioral roles reflect region-specific circuit wiring

The behavioral dissociation between iHipp and vHipp OLM^α2^ cells mirrors their anatomical segregation. Manipulating iHipp OLM^α2^ cells altered object approach behaviors but did not induce stable memory (Figures 2 and 3), consistent with studies showing that intermediate CA1 supports encoding and discrimination of object/place information (Siwani et al., 2018). Also, intermediate OLM^α2^ cells receive additional subicular and raphe inputs that could support object-related processing (Thulin et al. 2025). Conversely, inhibition of vHipp OLM^α2^ cells increased center avoidance, consistent with anxiogenic phenotypes, and activation modulated locomotor arousal, in line with the ventral hippocampus’ established role in emotional processing (Figures 2 and 3F; Mikulovic et al., 2018: Jimenez et al., 2018; Adhikari et al., 2010). Unexpectedly, stimulation of BLA→vHipp projections produced a preference for the light-paired object (Figure 5J), even though vHipp OLM^α2^ manipulations did not induce such preference, vHipp activation can be interpreted as anxiety modulating in the NOR and COP tests (Figures 2 and 5). This apparent contradiction likely reflects engagement of distinct pyramidal neuron subpopulations: BLA targeting pyramidal neurons may mediate approach- or salience-driven behaviors, while OLM^α2^-sensitive pyramidal neurons mediate avoidance or emotional suppression. Such microcircuit selectivity has been previously described in other hippocampal-amygdala interactions (Correia et al., 2016; Felix-Ortiz et al., 2013).

### A mechanistic model of OLM^α2^ function along the hippocampal axis

Integrating these data together with previously published work (Siwani et al., 2018, Mikulovic et al., 2018), we propose the following conceptual model: OLM^α2^ interneurons regulate the entorhinal, CA3 balance in CA1, shaping pyramidal ensemble output. The downstream targets of these pyramidal neurons differ along the longitudinal axis, creating region-specific behavioral consequences. In iHipp, OLM^α2^ cells influence pyramidal neurons projecting to sensory cortices, supporting exploration and object-based encoding. In vHipp, OLM^α2^ cells influence pyramidal neurons projecting to BLA and nAcc, shaping arousal, avoidance, and emotional salience. Although vHipp receives strong BLA input, OLM^α2^ interneurons themselves are not the direct recipients, emphasizing a disynaptic modulatory role rather than direct limbic gating.

Input differences directly to OLM^α2^ cells along the dorsoventral axis further suggest that such intermediate cells may integrate contextual or neuromodulatory signals, providing a substrate for memory-related computations that are different from the emotional/arousal functions of ventral OLMα2 cells (Thulin et al., 2025).

This framework is consistent with work suggesting that the hippocampus contains functionally tuned microcircuits embedded within position-dependent large-scale networks (Strange et al., 2014; Bannerman et al., 2004). Our findings refine this view, showing that functional specialization arises not only from pyramidal output patterns but also from interneuron-specific input architecture along the hippocampal axis.

### Implications for hippocampal circuit function and disease

Because vHipp dysregulation is implicated in anxiety disorders, PTSD, and depression (Tovote et al., 2015; Fanselow and Dong, 2010), identifying interneuron-specific contributions to limbic circuit modulation holds translational relevance. Our data suggest that selective modulation of OLM^α2^ interneurons, which avoids direct activation of BLA neurons, may represent a route for tuning emotional responses without inducing generalized arousal, a desirable property for therapeutic strategies. Conversely, modulation of iHipp OLM^α2^ cells may selectively influence memory encoding without unintended effects on anxiety, highlighting the functional precision available through longitudinally targeted interventions of dementia.

## Concluding remarks

This study establishes that the behavioral roles of OLM^α2^ interneurons are mechanistically dictated by their anatomical position within the longitudinal hippocampal circuit. By regulating pyramidal neuron outputs to either sensory or affective structures, OLM^α2^ cells contribute to distinct and region-specific computations, revealing a fine-grained functional architecture that underlies the dual mnemonic and emotional roles of the hippocampus.

## List of abbreviations

dHipp: Dorsal hippocampus
iHipp: Intermediate hippocampus
vHipp: Ventral hippocampus
OLM: Oriens lacunosum-moleculare
BLA: Basolateral amygdala
nAcc: Nucleus accumbens
NOR: Novel object recognition
COP: Conditioned odor preference
ChR2: Channelrhodopsin-2
Arch: Archaerhodopsin
POA: Predator odor aversion
eYFP: Enhanced yellow fluorescent protein
GFP: Green fluorescent protein
CamKII: Calmodulin-dependant Kinase II
AAV: Adeno-associated virus

## Acknowledgements

We thank Uppsala University Behavioral Facility (UUBF) for support and the Viral Core Facility (VCF, Charité, Berlin), and Ed Callaway for the production of the helper and pseudotyped rabies viruses. We thank the Swedish Research Council (2022-01245; www.vr.se) the Swedish Brain Foundation (FO2022-0018, PS2021-0061; http://hjarnfonden.se), Olle Engkvist Byggmästare Foundation (220-0254; https://engkviststiftelserna.se) and U-Share (SLU 2021.1.1.1-4394)

## Author contributions

This project was conceptualized by S.S. and K.K., and supervised by K.K. and E.R. Funding was acquired by S.S., K.K., and E.R. Resources and materials were obtained by K.K. Methodology and experimentation was conducted by S.S., A.T., A.F., A.L. Formal analysis was done by S.S. and visualization by S.S. and K.K. The original draft was written by S.S. and K.K., edited by S.S., K.K. and E.R and reviewed by all authors.

## Declaration of generative AI and AI-assisted technologies

Authors of this article used chatGPT to improve wording and grammar. The writing was reviewed and edited by the authors who take full responsibility for the contents of the publication.

## References

Adhikari, A., Topiwala, M. A., & Gordon, J. A. (2010). Synchronized activity between the ventral hippocampus and the medial prefrontal cortex during anxiety. Neuron. 2010 Jan 28;65(2):257–69. doi: 10.1016/j.neuron.2009.12.002.

Admon, R., Leykin, D., Lubin, G., Engert, V., Andrews, J., Pruessner, J., & Hendler, T. (2013). Stress-induced reduction in hippocampal volume and connectivity with the ventromedial prefrontal cortex are related to maladaptive responses to stressful military service. Human Brain Mapping, 34(11), 2808–2820. 10.1002/hbm.22100

Amaral, D.G., Witter, M.P., 1989. The three-dimensional organization of the hippocampal formation: A review of anatomical data. Neuroscience 31, 571–591. 10.1016/0306-4522(89)90424-7

Bast, T., Wilson, I.A., Witter, M.P., Morris, R.G.M., 2009. From Rapid Place Learning to Behavioral Performance: A Key Role for the Intermediate Hippocampus. PLoS Biol. 7, e1000089. 10.1371/journal.pbio.1000089

Bland, B.H., 1986. The physiology and pharmacology of hippocampal formation theta rhythms. Prog. Neurobiol. 26, 1–54. 10.1016/0301-0082(86)90019-5

Britt, J. P., Benaliouad, F., McDevitt, R. A., Stuber, G. D., Wise, R. A., & Bonci, A. (2012). Synaptic and behavioral profile of multiple glutamatergic inputs to the nucleus accumbens. Neuron, 76(4), 790–803. 10.1016/j.neuron.2012.09.040.

Burton, B.G., Hok, V., Save, E., Poucet, B., 2009. Lesion of the ventral and intermediate hippocampus abolishes anticipatory activity in the medial prefrontal cortex of the rat. Behav. Brain Res. 199, 222–234. 10.1016/j.bbr.2008.11.045

Cembrowski, M. S., Wang, L., Sugino, K., Shields, B. C., & Spruston, N. (2016). Hipposeq: A comprehensive RNA-seq database of gene expression in hippocampal principal neurons. eLife, 5, e14997. 10.7554/eLife.14997

Cholvin, T., Loureiro, M., Cassel, R., Cosquer, B., Herbeaux, K., de Vasconcelos, A.P., Cassel, J.-C., 2016. Dorsal hippocampus and medial prefrontal cortex each contribute to the retrieval of a recent spatial memory in rats. Brain Struct. Funct. 221, 91–102. 10.1007/s00429-014-0894-6

Crawley, J.N., Chen, T., Puri, A., Washburn, R., Sullivan, T.L., Hill, J.M., Young, N.B., Nadler, J.J., Moy, S.S., Young, L.J., Caldwell, H.K., Young, W.S., 2007. Social approach behaviors in oxytocin knockout mice: Comparison of two independent lines tested in different laboratory environments. Neuropeptides 41, 145–163. 10.1016/j.npep.2007.02.002

Dulawa, S.C., Grandy, D.K., Low, M.J., Paulus, M.P., Geyer, M.A., 1999. Dopamine D4 Receptor-Knock-Out Mice Exhibit Reduced Exploration of Novel Stimuli. J. Neurosci. 19, 9550–9556. 10.1523/JNEUROSCI.19-21-09550.1999

Fanselow, M.S., Dong, H.-W., 2010. Are the Dorsal and Ventral Hippocampus Functionally Distinct Structures? Neuron 65, 7–19. 10.1016/j.neuron.2009.11.031

Goodrich-Hunsaker, N.J., Hunsaker, M.R., Kesner, R.P., 2008. The interactions and dissociations of the dorsal hippocampus subregions: How the dentate gyrus, CA3, and CA1 process spatial information. Behav. Neurosci. 122, 16–26. 10.1037/0735-7044.122.1.16

Huff, M.L., Emmons, E.B., Narayanan, N.S., LaLumiere, R.T., 2016. Basolateral amygdala projections to ventral hippocampus modulate the consolidation of footshock, but not contextual, learning in rats. Learn. Mem. 23, 51–60. 10.1101/lm.039909.115

Hunsaker, M.R., Kesner, R.P., 2008. Evaluating the differential roles of the dorsal dentate gyrus, dorsal CA3, and dorsal CA1 during a temporal ordering for spatial locations task. Hippocampus 18, 955–964. 10.1002/hipo.20455

Hunsaker, M.R., Rosenberg, J.S., Kesner, R.P., 2008. The role of the dentate gyrus, CA3a,b, and CA3c for detecting spatial and environmental novelty. Hippocampus 18, 1064–1073. 10.1002/hipo.20464

Janak, P. H., & Tye, K. M. (2015). From circuits to behaviour in the amygdala. Nature, 517(7534), 284–292. 10.1038/nature14188

Kesner, R.P., Hunsaker, M.R., Ziegler, W., 2011. The role of the dorsal and ventral hippocampus in olfactory working memory. Neurobiol. Learn. Mem. 96, 361–366. 10.1016/j.nlm.2011.06.011

Kjelstrup, K. G., Tuvnes, F. A., Steffenach, H., Murison, R., Moser, E. I., & Moser, M. B. (2002). Reduced fear expression after lesions of the ventral hippocampus. Proceedings of the National Academy of Sciences of the United States of America, 99(16), 10825–10830. 10.1073/pnas.152112399

Kramis, R., Vanderwolf, C.H., Bland, B.H., 1975. Two types of hippocampal rhythmical slow activity in both the rabbit and the rat: Relations to behavior and effects of atropine, diethyl ether, urethane, and pentobarbital. Exp. Neurol. 49, 58–85. 10.1016/0014-4886(75)90195-8

Leão, R.N., Mikulovic, S., Leão, K.E., Munguba, H., Gezelius, H., Enjin, A., Patra, K., Eriksson, A., Loew, L.M., Tort, A.B.L., Kullander, K., 2012. OLM interneurons differentially modulate CA3 and entorhinal inputs to hippocampal CA1 neurons. Nat. Neurosci. 15, 1524–1530. 10.1038/nn.3235

LeGates, T.A., Kvarta, M.D., Tooley, J.R., Francis, T.C., Lobo, M.K., Creed, M.C., Thompson, S.M., 2018. Reward behaviour is regulated by the strength of hippocampus-nucleus accumbens synapses. Nature 564, 258–262. 10.1038/s41586-018-0740-8

Lovett-Barron, M., Kaifosh, P., Kheirbek, M.A., Danielson, N., Zaremba, J.D., Reardon, T.R., Turi, G.F., Hen, R., Zemelman, B.V., Losonczy, A., 2014. Dendritic Inhibition in the Hippocampus Supports Fear Learning. Science 343, 857–863. 10.1126/science.1247485

Mikulovic, S., Restrepo, C.E., Siwani, S., Bauer, P., Pupe, S., Tort, A.B.L., Kullander, K., Leão, R.N., 2018. Ventral hippocampal OLM cells control type 2 theta oscillations and response to predator odor. Nat. Commun. 9, 3638. 10.1038/s41467-018-05907-w

McHugh, S. B., Deacon, R. M. J., Rawlins, J. N. P., & Bannerman, D. M. (2004). Amygdala and ventral hippocampus contribute differentially to mechanisms of fear and anxiety. Behavioral Neuroscience, 118(1), 63–78. 10.1037/0735-7044.118.1.63

Montoya, C.P., Heynen, A.J., Faris, P.D., Sainsbury, R.S., 1989. Modality specific Type 2 theta production in the immobile rat. Behav. Neurosci. 103, 106–111. 10.1037/0735-7044.103.1.106

Moser, E.I., Kropff, E., Moser, M.-B., 2008. Place Cells, Grid Cells, and the Brain’s Spatial Representation System. Annu. Rev. Neurosci. 31, 69–89. 10.1146/annurev.neuro.31.061307.090723

Phillips, R.G., LeDoux, J.E., 1992. Differential contribution of amygdala and hippocampus to cued and contextual fear conditioning. Behav. Neurosci. 106, 274–285. 10.1037/0735-7044.106.2.274

Pi, G., Gao, D., Wu, D., Wang, Y., Lei, H., Zeng, W., Gao, Y., Yu, H., Xiong, R., Jiang, T., Li, S., Wang, X., Guo, J., Zhang, S., Yin, T., He, T., Ke, D., Li, R., Li, H., Liu, G., Yang, X., Luo, M., Zhang, X., Yang, Y., Wang, J., 2020. Posterior basolateral amygdala to ventral hippocampal CA1 drives approach behaviour to exert an anxiolytic effect. Nat. Commun. 11, 183. 10.1038/s41467-019-13919-3

Preibisch, S., Saalfeld, S., Tomancak, P., 2009. Globally optimal stitching of tiled 3D microscopic image acquisitions. Bioinformatics 25, 1463–1465. 10.1093/bioinformatics/btp184

Risold, P.Y., Swanson, L.W., 1996. Structural Evidence for Functional Domains in the Rat Hippocampus. Science 272, 1484–1486. 10.1126/science.272.5267.1484

Sanders, M.J., Wiltgen, B.J., Fanselow, M.S., 2003. The place of the hippocampus in fear conditioning. Eur. J. Pharmacol. 463, 217–223. 10.1016/S0014-2999(03)01283-4

Scoville, W.B., Milner, B., 1957. LOSS OF RECENT MEMORY AFTER BILATERAL HIPPOCAMPAL LESIONS. J. Neurol. Neurosurg. Psychiatry 20, 11–21. 10.1136/jnnp.20.1.11

Siwani, S., França, A.S.C., Mikulovic, S., Reis, A., Hilscher, M.M., Edwards, S.J., Leão, R.N., Tort, A.B.L., Kullander, K., 2018. OLMα2 Cells Bidirectionally Modulate Learning. Neuron 99, 404–412.e3. 10.1016/j.neuron.2018.06.022

Staples, L.G., 2010. Predator odor avoidance as a rodent model of anxiety: Learning-mediated consequences beyond the initial exposure. Neurobiol. Learn. Mem. 94, 435–445. 10.1016/j.nlm.2010.09.009

Strange, B.A., Witter, M.P., Lein, E.S., Moser, E.I., 2014. Functional organization of the hippocampal longitudinal axis. Nat. Rev. Neurosci. 15, 655–669. 10.1038/nrn3785

Tervo, D.G.R., Hwang, B.-Y., Viswanathan, S., Gaj, T., Lavzin, M., Ritola, K.D., Lindo, S., Michael, S., Kuleshova, E., Ojala, D., Huang, C.-C., Gerfen, C.R., Schiller, J., Dudman, J.T., Hantman, A.W., Looger, L.L., Schaffer, D.V., Karpova, A.Y., 2016. A Designer AAV Variant Permits Efficient Retrograde Access to Projection Neurons. Neuron 92, 372–382. 10.1016/j.neuron.2016.09.021

Thulin A, Henriksson K, Nogueira I, Kullander K. Differentiated Presynaptic Input to OLMα2 Cells Along the Hippocampal Dorsoventral Axis: Implications for Hippocampal Microcircuit Function. Hippocampus. 2025 Sep;35(5):e70026. doi: 10.1002/hipo.70026. PMID: 40757734; PMCID: PMC12320472.

Toledo-Rodriguez, M., Sandi, C., 2011. Stress during Adolescence Increases Novelty Seeking and Risk-Taking Behavior in Male and Female Rats. Front. Behav. Neurosci. 5. 10.3389/fnbeh.2011.00017

Wang, M.E., Fraize, N.P., Yin, L., Yuan, R.K., Petsagourakis, D., Wann, E.G., Muzzio, I.A., 2013. Differential roles of the dorsal and ventral hippocampus in predator odor contextual fear conditioning. Hippocampus 23, 451–466. 10.1002/hipo.22105

Wersinger, S.R., Caldwell, H.K., Martinez, L., Gold, P., Hu, S.-B., Young, W.S., 2007. Vasopressin 1a receptor knockout mice have a subtle olfactory deficit but normal aggression. Genes Brain Behav. 6, 540–551. 10.1111/j.1601-183X.2006.00281.x

Woodley, S.K., Baum, M.J., 2003. Effects of sex hormones and gender on attraction thresholds for volatile anal scent gland odors in ferrets. Horm. Behav. 44, 110–118. 10.1016/S0018-506X(03)00126-0

Wrenn, C.C., Kinney, J.W., Marriott, L.K., Holmes, A., Harris, A.P., Saavedra, M.C., Starosta, G., Innerfield, C.E., Jacoby, A.S., Shine, J., Iismaa, T.P., Wenk, G.L., Crawley, J.N., 2004. Learning and memory performance in mice lacking the GAL-R1 subtype of galanin receptor. Eur. J. Neurosci. 19, 1384–1396. 10.1111/j.1460-9568.2004.03214.x

Xu, F., Ono, M., Ito, T., Uchiumi, O., Wang, F., Zhang, Y., Sun, P., Zhang, Q., Yamaki, S., Yamamoto, R., Kato, N., 2020. Remodeling of projections from ventral hippocampus to prefrontal cortex in Alzheimer’s mice. J. Comp. Neurol. cne.25032. 10.1002/cne.25032

Yang, M., Crawley, J.N., 2009. Simple Behavioral Assessment of Mouse Olfaction. Curr. Protoc. Neurosci. 48. 10.1002/0471142301.ns0824s48

Yang, Y., Wang, J.-Z., 2017. From Structure to Behavior in Basolateral Amygdala-Hippocampus Circuits. Front. Neural Circuits 11, 86. 10.3389/fncir.2017.00086

