## Supplementary material for "Conserved inhibitory motifs yield distinct hippocampal functions through regional circuit embedding"

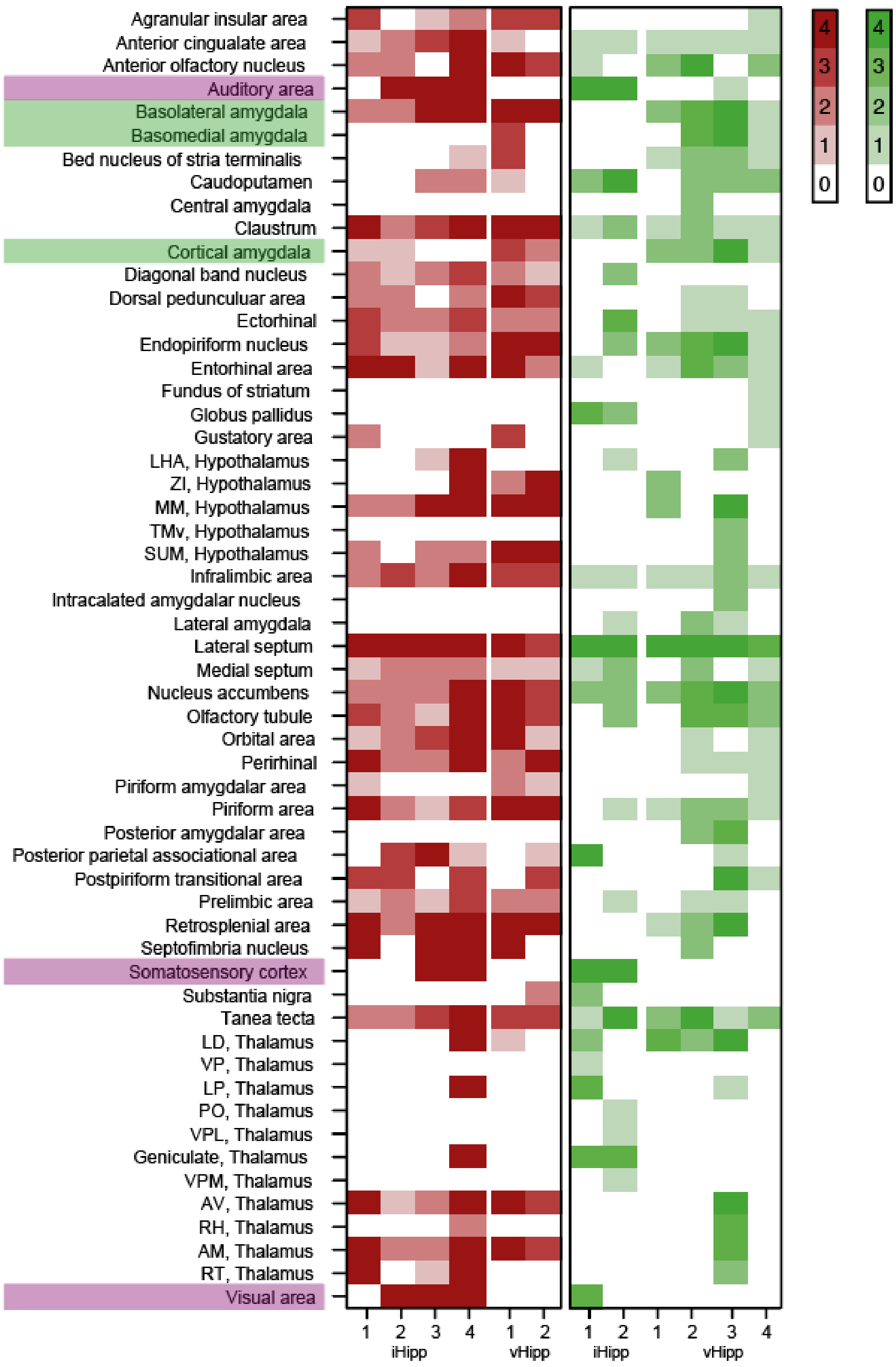


**Supplementary Figure 1.** Color-coded heatmap showing projection strength across individual structures and mice (Each column represents an individual animal). Comparisons are shown for iHipp (left) and vHipp (right), with expression intensity represented on a five-point scale (0–4). Red represent tracings performed with the retrograde rAAV-CAG-hChR2(H134R)-tdTomato virus and green with the rAAV-CAG-hChR2(H134R)-tdTomato virus as an anterograde tracer. Highlighted in green and purple are structures that potentially have explanatory power in the observed behavioral differences comparing iHipp (purple) and vHipp (green).

***Supplementary Table 1.*** *Materials*

| Viruses | | | |
|---|---|---|---|
| Virus | **Titer** | **Source** | **Identifier** |
| AAV2-Ef1a-DIO-hChR2(H134R)-EYFP | 4x10^12vg/ml | UNC vector core | LOT: av4278p |
| AAV5-CamKIIa-eArch3-eYFP | | | LOT: 4840B |
| (Retro) AAV-CAG-tdTomato | 4.8x 10e12 vg/ml | UNC vector core | LOT: 116644 |
| (Retro) AAV-CAG-CRE | 1,50E+12 vg/ml | UNC vector core | Lot: AV7703H |
| (Retro) AAV-CAG-hChR2-H134-tdTomato | 7,60E+12 GC/ml | Addgene | Lot: v38830 |
| AAV9-Flex-Arch-GFP | 2,00E+12 gc/ml | Addgene | Lot: 117525 |
| AAV2-EF1a-DIO-Arch3.0-eYFp | | | Lot: AV4881c, AV4881D |
| (Retro) AAV-Syn-H2B-GFP-TVA-N2c.G-WPRE3 | 8,69E+11 VG/ml | Charité | Virus no: BA-335d |
| RABV CVS-N2c(deltaG)-mCherry) | 2.86E+10 IU/ml | Charité | Lot: BRVena-32a |

| Optical equipment | | |
|---|---|---|
| Material | **Model/type** | **Source** |
| Fiber optic cannula | MFC_200/230-0.37_4mm_ZF1.25(G)_FLT | Doric lenses |
| Rotator | | Doric lenses |
| Fiberoptic patch cord | 2 SBP(2)_200/240/1100-0.22_1.5m_FCM-2xZF1.25(FP) | Doric lenses |
| Laser, green 535 nm | | Shangai Dream Lasers |

| | | |
|---|---|---|
| Larser, Blue 473 nm | | Shangai Dream Lasers |
| DAQ box | | National instruments |
| Zeiss BF microscope | Imager .Z2 | Zeiss |
| Olympus BF microscope | BX61WI | Olympus |

| Surgical equipment | | |
|---|---|---|
| Material | **Model/Type** | **Source** |
| Anchor screws | MCS1x2, HM71009 | Agnthos |
| Dental cement primary + primer | Optibond FL | MW dental |
| Dental cement secondary | Tetric evoflow Syringe A1 | MW dental |

| Software | | |
|---|---|---|
| Program | **Version** | **Source** |
| Ethovision | XT14-16 | Noldus |
| LabVIEW NXG Full Development System | Custom | National instruments |
| Graphad prism | Version 10 | GraphPad software |
| Biorender | | Biorender |
| ImageJ | (Plugin: Cell counter (PRID:SCR_025376)) | Fiji |

| Reagents (IHC) | | |
|---|---|---|
| Type | **Source** | **Identifier** |
| Antibody, Rabbit anti-cFOS (9F6) | Cell Signaling | #2250 |
| Antibody, donkey anti-rabbit 647 | Invitrogen | A31573, Lot#2674379 |
| Antibody, DAPI | VWR | CatNo.: 40011 |
| Donkey Serum | Biowest | S2170-100 |
| BSA | Sigma-Aldrich | A4503 |
| Triton-X-100 | Sigma-Aldrich | T9284-500ML |
| Tween 20 | Sigma-Aldrich | P1379-500ML |
| Gelatine | Fisher Scientific | G/0150/53 |
| PBS | ThermoFisher | CatNo.: 10051163 |
| Fluoroshield | Abcam | ab104135 |

| Drugs and other chemicals | | | |
|---|---|---|---|
| Type (active substance) | **Concentration** | **Source** | **Identifier** |
| Ketalar (Ketamine) | 10mg/mL and 50 mg/mL | Apoteket | 325008 |
| Domitor (Medetomidinhydrochloride) | 1mg/mL | Apoteket | Vnr: 569830 |
| Vetergesic (Buprenorphine) | 0.3mg/ml | Apoteket | Vnr: 054594 |

| | | | |
|---|---|---|---|
| Isoflurane Baxter | | Apoteket | Vnr: 476843 |
| Rimadyl bovis vet. (carprofen) | 50mg/ml | Apoteket | p2802-05 |
| Marcain (bupivakain) | 2.5mg/ml | Apoteket | 9953745-1 |
| Klorhexidin | 0.5mg/ml | Apoteket | Vnr: 537977 |
| Eyegel, Oftagel | 2.5mg/g | Apoteket | Vnr: 404426 |
| Vanillin | | Sigma-Aldrich | Product nr: W310727 |
| Isoamyl alcohol | | Sigma-Aldrich | Product nr: W205710 |

***Supplementary Table 2.*** *Descriptive statistics Figure 2 and 3*

| Figure | Group | N | Minimum | 25% | Median | 75% | Maximum | Mean | Sd | SEM |
|---|---|---|---|---|---|---|---|---|---|---|
| 2D | Control | 12 | 21,48 | 35,41 | 62,14 | 177,8 | 221,8 | 94,58 | 75,86 | 21,9 |
| | $OLM^{\alpha2\text{-}Arch}$ | 13 | 10,2 | 26,4 | 45,68 | 155,2 | 235,7 | 83,14 | 77,25 | 21,42 |
| | Control | 12 | 2,8 | 11,41 | 15,4 | 46,35 | 143,6 | 32,88 | 40,37 | 11,65 |
| | $OLM^{\alpha2\text{-}Arch}$ | 13 | 0,24 | 6 | 12,64 | 65,6 | 143,2 | 38,5 | 46,26 | 12,83 |
| 2E | Control | 9 | 0,5844 | 0,6767 | 1,117 | 1,858 | 2,908 | 1,317 | 0,8033 | 0,284 |
| | $OLM^{\alpha2\text{-}ChR2}$ | 9 | 0,2946 | 1,03 | 1,477 | 1,719 | 1,875 | 1,354 | 0,4777 | 0,1511 |
| 2G | Control | 9 | 0,5844 | 0,6767 | 1,117 | 1,858 | 2,908 | 1,317 | 0,8033 | 0,284 |
| | $OLM^{\alpha2\text{-}ChR2}$ | 9 | 0,2946 | 1,03 | 1,477 | 1,719 | 1,875 | 1,354 | 0,4777 | 0,1511 |
| | Control | 7 | 0,8296 | 0,9468 | 1,144 | 1,815 | 1,921 | 1,325 | 0,4475 | 0,1691 |
| | $OLM^{\alpha2\text{-}Arch}$ | 7 | 0,6164 | 0,9718 | 1,768 | 1,856 | 2,94 | 1,576 | 0,7708 | 0,2913 |
| 2H | Control | 9 | 1266 | 1558 | 2003 | 2351 | 2762 | 1982 | 486,8 | 162,3 |
| | $OLM^{\alpha2\text{-}ChR2}$ | 9 | 1491 | 2524 | 3008 | 3161 | 3386 | 2799 | 571,2 | 190,4 |
| | Control | 7 | 2069 | 2243 | 2794 | 2842 | 3026 | 2579 | 376,8 | 142,4 |
| | $OLM^{\alpha2\text{-}Arch}$ | 7 | 747,9 | 1178 | 1860 | 2493 | 2685 | 1819 | 705,7 | 266,7 |
| 3C | Control | 10 | 2022 | 2594 | 4320 | 6182 | 7154 | 4452 | 1923 | 608.2 |

| | | | | | | | | | | |
|---|---|---|---|---|---|---|---|---|---|---|
| | OLM$^{\alpha2\text{-}ChR2}$ | 9 | 2543 | 3736 | 5469 | 5971 | 7141 | 5069 | 1472 | 490.8 |
| | Control | 12 | 1564 | 1920 | 2107 | 3030 | 4402 | 2518 | 870.9 | 251.4 |
| | OLM$^{\alpha2\text{-}Arch}$ | 8 | 1745 | 1985 | 2286 | 4109 | 5022 | 2856 | 1235 | 436.7 |
| 3D | Control | 9 | 2314 | 2464 | 2635 | 3143 | 4018 | 2855 | 535.8 | 178.6 |
| | OLM$^{\alpha2\text{-}ChR2}$ | 9 | 1560 | 2492 | 3286 | 4080 | 4308 | 3195 | 914.7 | 304.9 |
| | Control | 7 | 1411 | 2259 | 2766 | 3106 | 3663 | 2675 | 714.3 | 270 |
| | OLM$^{\alpha2\text{-}Arch}$ | 7 | 894.8 | 1155 | 1405 | 3220 | 4626 | 2031 | 1377 | 520.4 |
| 3E | Control | 10 | 6.04 | 24.55 | 43.7 | 60.5 | 115.9 | 46.25 | 31.38 | 9.922 |
| | OLM$^{\alpha2\text{-}ChR2}$ | 9 | 24.32 | 30.4 | 52.6 | 87.84 | 127.6 | 58.8 | 37.35 | 12.45 |
| | Control | 12 | 4 | 9.01 | 23.18 | 55.87 | 71.48 | 32.96 | 25.53 | 7.371 |
| | OLM$^{\alpha2\text{-}Arch}$ | 8 | 12.72 | 14.83 | 35.96 | 49.87 | 58.44 | 33.38 | 17.59 | 6.219 |
| 3F | Control | 9 | 20.24 | 73.38 | 170 | 217 | 244 | 147.6 | 78.25 | 26.08 |
| | OLM$^{\alpha2\text{-}ChR2}$ | 9 | 14.56 | 91.62 | 217.4 | 254.1 | 271.2 | 173.9 | 93.04 | 31.01 |
| | Control | 7 | 20.12 | 37.39 | 116.9 | 204.2 | 248.1 | 125.7 | 86.77 | 30.68 |
| | OLM$^{\alpha2\text{-}Arch}$ | 7 | 12.84 | 22.92 | 42.8 | 71.12 | 71.76 | 45.35 | 24.84 | 9.39 |
| 3G | Control | 10 | 22.33 | 39.07 | 47.79 | 58.39 | 66.51 | 47.42 | 14.06 | 4.446 |
| | | 10 | 33.49 | 41.61 | 52.21 | 60.93 | 77.67 | 52.58 | 14.06 | 4.446 |
| | OLM$^{\alpha2\text{-}ChR2}$ | 9 | 43.83 | 52.89 | 63.06 | 74.02 | 89.45 | 63.18 | 14.43 | 4.809 |
| | | 9 | 10.55 | 25.98 | 36.94 | 47.11 | 56.17 | 36.82 | 14.43 | 4.809 |
| | Control | 12 | 32.36 | 41.66 | 56.9 | 62.37 | 78.25 | 54.82 | 14.11 | 4.072 |
| | | 12 | 21.75 | 37.63 | 43.1 | 58.34 | 67.64 | 45.18 | 14.11 | 4.072 |
| | OLM$^{\alpha2\text{-}Arch}$ | 8 | 24.19 | 32.2 | 41.68 | 53.25 | 63.39 | 42.7 | 12.78 | 4.519 |
| | | 8 | 36.61 | 46.75 | 58.32 | 67.8 | 75.81 | 57.3 | 12.78 | 4.519 |
| 3H | Control | 9 | 35.23 | 41.38 | 45.99 | 58.85 | 69.01 | 49.59 | 11.33 | 3.777 |
| | | 9 | 30.99 | 41.15 | 54.01 | 58.62 | 64.77 | 50.41 | 11.33 | 3.777 |
| | OLM$^{\alpha2\text{-}ChR2}$ | 9 | 37.05 | 46.03 | 51.64 | 53.4 | 61.76 | 50.17 | 6.797 | 2.266 |
| | | 9 | 38.24 | 46.6 | 48.36 | 53.97 | 62.95 | 49.83 | 6.797 | 2.266 |
| | Control | 7 | 14.87 | 30.65 | 43.36 | 56.76 | 72.83 | 43.46 | 18.61 | 7.035 |
| | | 7 | 27.17 | 43.24 | 56.64 | 69.35 | 85.13 | 56.54 | 18.61 | 7.035 |
| | OLM$^{\alpha2\text{-}Arch}$ | 7 | 36.7 | 49.38 | 56.7 | 65.52 | 65.86 | 54.72 | 10.38 | 3.923 |
| | | 7 | 34.14 | 34.48 | 43.3 | 50.62 | 63.3 | 45.28 | 10.38 | 3.923 |

***Supplementary Table 3***. *Test results Figures 2 and 3*

| Figure | Groups | n | Test | Results | |
|---|---|---|---|---|---|
| 2D | Control | 12 | Mann-Whitney | P value | 0.4371 |
| | OLM$^{\alpha2\text{-}Arch}$ | 13 | | U | 63 |
| | Control | 12 | Mann-Whitney | P value | 0.7167 |
| | OLM$^{\alpha2\text{-}Arch}$ | 13 | | U | 71 |
| 2E | Control | 12 | Mann-Whitney | P value | 0.9049 |
| | OLM$^{\alpha2\text{-}Arch}$ | 13 | | U | t=0,1208, df=23 |
| 2G | Control | 9 | Unpaired T-test | P value | 0.8918 |
| | OLM$^{\alpha2\text{-}ChR2}$ | 9 | | t, df | t=0,1382, df=16 |
| | Control | 7 | Unpaired T-test | P value | 0.4691 |
| | OLM$^{\alpha2\text{-}Arch}$ | 7 | | t, df | t=0,7477, df=12 |
| 2H | Control | 9 | Unpaired T-test | P value | 0.0049** |
| | OLM$^{\alpha2\text{-}ChR2}$ | 9 | | t, df | t=3,266, df=16 |
| | Control | 7 | Unpaired T-test | P value | 0.0436* |
| | OLM$^{\alpha2\text{-}Arch}$ | 7 | | t, df | t=2,235, df=13 |
| 3C | Control | 10 | Unpaired T-test | P value | 0.4472 |
| | OLM$^{\alpha2\text{-}ChR2}$ | 9 | | t, df | t=0,7780, df=17 |
| | Control | 12 | Mann-Whitney | P value | 0.5208 |
| | OLM$^{\alpha2\text{-}Arch}$ | 8 | | U | 39 |
| 3D | Control | 9 | Unpaired T-test | P value | 0.3508 |
| | OLM$^{\alpha2\text{-}ChR2}$ | 9 | | t, df | t=0,9611, df=16 |
| | Control | 7 | Mann-Whitney | P value | 0.2086 |
| | OLM$^{\alpha2\text{-}Arch}$ | 7 | | U | 14 |
| 3E | Control | 10 | Mann-Whitney | P value | 0.9048 |
| | OLM$^{\alpha2\text{-}ChR2}$ | 9 | | U | 43 |
| | Control | 12 | Unpaired T-test | P value | 0.7276 |
| | OLM$^{\alpha2\text{-}Arch}$ | 8 | | t, df | t=0,3538, df=18 |
| 3F | Control | 9 | Unpaired T-test | P value | 0.5252 |
| | OLM$^{\alpha2\text{-}ChR2}$ | 9 | | t, df | t=0,6496, df=16 |
| | Control | 7 | Unpaired T-test | P value | 0.0138* |
| | OLM$^{\alpha2\text{-}Arch}$ | 7 | | t, df | t=2,882, df=12 |

| | | | | | |
|---|---|---|---|---|---|
| 3G | Control | 10 | Paired T-test | P value | 0.5766 |
| | | | | t, df | t=0,5794, df=9 |
| | OLM$^{\alpha2\text{-}ChR2}$ | 9 | Paired T-test | P value | 0.0276* |
| | | | | t, df | t=2,410, df=17 |
| | Control | 10 | Unpaired T-test | P value | 0.0254* |
| | OLM$^{\alpha2\text{-}ChR2}$ | 9 | | t, df | t=2,741, df=8 |
| | Control | 12 | Paired T-test | P value | 0.2617 |
| | | | | t, df | t=1,183, df=11 |
| | OLM$^{\alpha2\text{-}Arch}$ | 8 | Paired T-test | P value | 0.1503 |
| | | | | t, df | t=1,615, df=7 |
| 3H | Control | 9 | Paired T-test | P value | 0.9165 |
| | | | | t, df | t=0,1082, df=8 |
| | OLM$^{ChR2}$ | 9 | Paired T-test | P value | 0.9406 |
| | | | | t, df | t=0,07692, df=8 |
| | Control | 7 | Paired T-test | P value | 0.3884 |
| | | | | t, df | t=0,9297, df=6 |
| | OLM$^{\alpha2\text{-}Arch}$ | 7 | Paired T-test | P value | 0.2746 |
| | | | | t, df | t=1,202, df=6 |

***Supplementary Table 4.*** *Descriptive statistics Figures 5 and 7*

| Figure | Group | N | Minimum | 25% | Median | 75% | Maximum | Mean | Sd | SEM |
|---|---|---|---|---|---|---|---|---|---|---|
| 5E | Left | 7 | 117 | 131 | 157 | 181 | 1151 | 295.6 | 377.9 | 142.8 |
| | Right | 7 | 150 | 153 | 231 | 326 | 1392 | 381.4 | 449.7 | 170 |
| 5H | Control | 7 | -152 | -90 | -16 | 53 | 89 | -17.86 | 82.34 | 31.12 |

| | | | | | | | | | | |
|---|---|---|---|---|---|---|---|---|---|---|
| | BLA$^{ChR2}$ | 7 | 22 | 26 | 55 | 145 | 241 | 85.86 | 80.64 | 30.48 |
| 5J | Control | 7 | 42.83 | 43.72 | 49.61 | 63.91 | 71.29 | 53.18 | 10.6 | 4.008 |
| | Control | 7 | 28.71 | 36.09 | 50.39 | 56.28 | 57.17 | 46.82 | 10.6 | 4.008 |
| | BLA$^{ChR2}$ | 7 | 50.51 | 51.69 | 54.35 | 66.84 | 72.95 | 57.96 | 8.553 | 3.233 |
| | BLA$^{ChR2}$ | 7 | 27.05 | 33.16 | 45.65 | 48.31 | 49.49 | 42.04 | 8.553 | 3.233 |
| 5K | Control | 7 | 25.32 | 72.16 | 132.7 | 193.8 | 223.3 | 125.8 | 71.93 | 27.19 |
| | BLA$^{ChR2}$ | 7 | 0 | 64.88 | 185.3 | 200.2 | 249.3 | 148.5 | 86.32 | 32.63 |
| 5L | Control | 7 | 1990 | 2585 | 3429 | 4579 | 4788 | 3512 | 1048 | 396.1 |
| | BLA$^{ChR2}$ | 7 | 2465 | 3672 | 4164 | 5049 | 5768 | 4293 | 1076 | 406.7 |
| 5N | Control | 7 | -1.195 | -0.3037 | -0.1135 | 0.147 | 1.976 | 0.0687 | 0.9538 | 0.3605 |
| | BLA$^{ChR2}$ | 8 | -1.565 | -0.9107 | -0.2097 | 1.207 | 2.663 | 0.07598 | 1.399 | 0.4945 |
| 5O | Control | 7 | -0.4202 | -0.3372 | -0.101 | 0.06588 | 0.2264 | -0.1221 | 0.2393 | 0.09045 |
| | BLA$^{ChR2}$ | 8 | -0.6296 | -0.2139 | -0.03323 | 0.472 | 0.668 | 0.02738 | 0.4325 | 0.1529 |
| 5P | Control | 7 | 1987 | 2196 | 2461 | 3031 | 3139 | 2544 | 425.7 | 160.9 |
| | BLA$^{ChR2}$ | 8 | 2149 | 2282 | 2400 | 2734 | 3021 | 2478 | 296.3 | 104.8 |
| | Control | 7 | 1340 | 1417 | 1740 | 2043 | 2114 | 1715 | 292.6 | 110.6 |
| | BLA$^{ChR2}$ | 8 | 1378 | 1425 | 1619 | 1970 | 2662 | 1746 | 427.5 | 151.1 |
| 5Q | Control | 7 | -746.1 | -394.8 | 73.7 | 558.4 | 607.5 | 19.94 | 496.8 | 187.8 |
| | BLA$^{ChR2}$ | 7 | 232.4 | 553.5 | 646.4 | 910.9 | 1023 | 665.6 | 255.2 | 96.45 |
| 7B | Control | 8 | -0.2407 | -0.131 | -0.0569 | 0.004118 | 0.2369 | -0.04263 | 0.1385 | 0.04897 |
| | OLM$^{\alpha2\text{-}Arch}$ | 7 | -0.3257 | -0.2423 | -0.1681 | 0.1225 | 0.1698 | -0.1064 | 0.1855 | 0.07013 |
| | Control | 9 | -0.7848 | 0.02847 | 0.05602 | 0.321 | 1.69 | 0.2007 | 0.6543 | 0.2181 |
| | OLM$^{\alpha2\text{-}ChR2}$ | 9 | -1.691 | -0.1264 | 0.08503 | 0.4635 | 0.9277 | 0.0139 | 0.7314 | 0.2438 |
| 7C | Control | 8 | -0.1699 | -0.103 | -0.01247 | 0.05967 | 0.07568 | -0.02198 | 0.08788 | 0.03107 |
| | OLM$^{\alpha2\text{-}Arch}$ | 7 | -0.1636 | -0.113 | -0.07887 | 0.04123 | 0.05467 | -0.0482 | 0.08592 | 0.03247 |
| | Control | 9 | -0.4402 | -0.2374 | 0.02069 | 0.3467 | 0.6955 | 0.05367 | 0.3813 | 0.1271 |
| | OLM$^{\alpha2\text{-}ChR2}$ | 8 | -0.06481 | 0.04527 | 0.4256 | 0.5609 | 0.8594 | 0.3765 | 0.3111 | 0.11 |
| 7D | Control | 8 | 1760 | 1822 | 1971 | 2109 | 2157 | 1968 | 144 | 50.91 |
| | OLM$^{\alpha2\text{-}Arch}$ | 7 | 1683 | 1728 | 1902 | 2264 | 2276 | 1973 | 262.1 | 99.07 |
| | Control | 8 | 1457 | 1529 | 1676 | 1822 | 1981 | 1685 | 174.6 | 61.74 |
| | OLM$^{\alpha2\text{-}Arch}$ | 7 | 1199 | 1369 | 1526 | 1917 | 1941 | 1617 | 298 | 112.6 |
| 7E | Control | 9 | 1638 | 1712 | 2181 | 2516 | 2763 | 2151 | 407.8 | 135.9 |
| | OLM$^{\alpha2\text{-}ChR2}$ | 9 | 1798 | 2114 | 2409 | 2765 | 3103 | 2426 | 422.9 | 141 |

| | | | | | | | | | | |
|---|---|---|---|---|---|---|---|---|---|---|
| | Control | 9 | 1126 | 1393 | 1770 | 2078 | 2318 | 1738 | 398.8 | 132.9 |
| | OLM$^{α2\text{-}ChR2}$ | 9 | 1366 | 1595 | 1849 | 2305 | 2677 | 1934 | 451 | 150.3 |
| 7F | Control | 9 | -696.1 | -487.9 | -79.21 | 385.2 | 650.3 | -52.92 | 462.4 | 154.1 |
| | OLM$^{α2\text{-}ChR2}$ | 9 | -741.2 | -673.9 | -293.6 | 609.1 | 968.9 | -77.87 | 654.5 | 218.2 |

***Supplementary Table 5.*** *Test results Figures 5 and 7*

| Figure | Groups | n | Test | Results | |
|---|---|---|---|---|---|
| 5E | Left BLA$^{ChR2}$ | 7 | Paired T-test | P value | 0.0347 |
| | Right BLA$^{ChR2}$ | 7 | | t, df | t=2,381, df=12 |
| 5H | Control | 7 | Wilcoxon | P value | 0.0156 * |
| | BLA$^{ChR2}$ | 7 | matched-pairs | W | 28 |
| 5J | Control | 7 | Paired T-test | P value | 0.4581 |
| | | | | t, df | t=0,7927, df=6 |
| | BLA$^{ChR2}$ | 7 | Paired T-test | P value | 0.049 * |
| | | | | t, df | t=2,462, df=6 |
| 5K | Control | 7 | Unpaired T-test | P value | 0.6026 |
| | BLA$^{ChR2}$ | 7 | | t, df | t=0,5347, df=12 |
| 5L | Control | 7 | Unpaired T-test | P value | 0.1937 |
| | BLA$^{ChR2}$ | 7 | | t, df | t=1,377, df=12 |
| 5N | Control | 7 | One sample T-test | P value | 0.8552 |
| | | | | t, df | t=0,1906, df=6 |
| | BLA$^{ChR2}$ | 8 | One sample T-test | P value | 0.8822 |
| | | | | t, df | t=0,1537, df=7 |
| 5O | Control | 7 | One sample T-test | P value | 0.2258 |
| | | | | t, df | t=1,350, df=6 |
| | BLA$^{ChR2}$ | 7 | One sample T-test | P value | 0.863 |
| | | | | t, df | t=0,1791, df=7 |
| 5Q | Control | 7 | One sample T-test | P value | 0.0099** |
| | BLA$^{ChR2}$ | 7 | One sample T-test | t, df | t=3,059, df=12 |

| | | | | | |
|---|---|---|---|---|---|
| 7B | Control | 8 | One sample T-test | P value | 0.4128 |
| | | | | t, df | t=0,8707, df=7 |
| | OLM$^{α2\text{-}Arch}$ | 7 | One sample T-test | P value | 0.1798 |
| | | | | t, df | t=1,518, df=6 |
| | Control | 9 | One sample Wilcox. | P value | 0.0977 |
| | | | | W | 29 |
| | OLM$^{α2\text{-}ChR2}$ | 9 | One sample T-test | P value | 0.9559 |
| | | | | t, df | t=0,05703, df=8 |
| 7C | Control | 8 | One sample T-test | P value | 0.5021 |
| | | | | t, df | t=0,7075, df=7 |
| | OLM$^{α2\text{-}Arch}$ | 7 | One sample T-test | P value | 0.1883 |
| | | | | t, df | t=1,484, df=6 |
| | Control | 9 | One sample T-test | P value | 0.6839 |
| | | | | t, df | t=0,4223, df=8 |
| | OLM$^{α2\text{-}ChR2}$ | 8 | One sample T-test | P value | 0.0111* |
| | | | | t, df | t=3,423, df=7 |
| 7F | Control | 9 | Unpaired T-test | P value | 0.9267 |
| | OLM$^{α2\text{-}ChR2}$ | 9 | | t, df | t=0,09343, df=16 |
| 7E | Left BLA$^{ChR2}$ | 7 | Paired T-test | P value | 0.0347 * |
| | Right BLA$^{ChR2}$ | 7 | | t, df | t=2,381, df=12 |
| 7H | Control | 7 | Wilcoxon | P value | 0.0156 * |
| | BLA$^{ChR2}$ | 7 | matched-pairs | W | 28 |

***Supplementary Table 6.*** *Test results Anova Figures 6 and 7*

| Figure | Groups | n | ANOVA table | SS | DF | MS | F (DFn, DFd) | P value |
|---|---|---|---|---|---|---|---|---|
| 5P | Control | 8 | Time x Treatment | 17353 | 1 | 17353 | F (1, 13) = 0,1816 | P=0,6770 |
| | | | Time | 4551785 | 1 | 4551785 | F (1, 13) = 47,63 | P<0,0001**** |
| | BLA$^{ChR2}$ | 7 | Treatment | 2337 | 1 | 2337 | F (1, 13) = 0,0134 8 | P=0,9093 |

| | | | | | | | | |
|---|---|---|---|---|---|---|---|---|
| | | | Subject | 2252788 | 13 | 173291 | F (13, 13) = 1,813 | P=0,1480 |
| 7D | Control | 7 | Time x Treatment | 9925 | 1 | 9925 | F (1, 13) = 0,2139 | P=0,6514 |
| | | | Time | 763720 | 1 | 763720 | F (1, 13) = 16,46 | P=0,0014** |
| | OLM$^{α2\text{-}Arch}$ | 8 | Treatment | 7199 | 1 | 7199 | F (1, 13) = 0,1336 | P=0,7206 |
| | | | Subject | 700368 | 13 | 53874 | F (13, 13) = 1,161 | P=0,3960 |
| 7E | Control | 9 | Time x Treatment | 14071 | 1 | 14071 | F (1, 16) = 0,2021 | P=0,6591 |
| | | | Time | 1843476 | 1 | 1843476 | F (1, 16) = 26,48 | P<0,0001**** |
| | OLM$^{α2\text{-}ChR2}$ | 9 | Treatment | 497072 | 1 | 497072 | F (1, 16) = 1,749 | P=0,2046 |
| | | | Subject | 4546491 | 16 | 284156 | F (16, 16) = 4,081 | P=0,0038 |
